# PRMT5 maintains tumor stem cells to promote pediatric high-grade glioma tumorigenesis

**DOI:** 10.1101/2024.03.12.583794

**Authors:** John DeSisto, Ilango Balakrishnan, Aaron J. Knox, Gabrielle Link, Sujatha Venkataraman, Rajeev Vibhakar, Adam L. Green

## Abstract

**Background:** Pediatric high-grade gliomas (PHGG) are aggressive, undifferentiated CNS tumors with poor outcomes, for which no standard-of-care drug therapy currently exists. Through a screen for epigenetic regulators, we identified *PRMT5* as essential for PHGG growth. We hypothesized that, similar to its effect in normal cells, PRMT5 promotes self-renewal of stem-like PHGG tumor initiating cells (TICs) essential for tumor growth. *Methods*. We conducted *in vitro* assays, including limiting dilution studies of self-renewal, to determine the phenotypic effects of *PRMT5* KD. We performed ChIP-Seq to identify PRMT5-mediated epigenetic changes and gene set enrichment analysis to identify pathways that PRMT5 regulates. Using an orthotopic xenograft model of PHGG, we tracked survival and histological characteristics resulting from *PRMT5* KD or administration of a PRMT5 inhibitor ± radiation therapy (RT).

**Results:** *In vitro*, *PRMT5* KD slowed cell cycle progression, tumor growth and self-renewal. *PRMT5* KD reduced H3K4me3 occupancy at genes associated with self-renewal, tumor formation and growth. *In vivo*, *PRMT5* KD increased survival and reduced tumor aggressiveness; however, pharmacological inhibition of PRMT5 with or without RT did not improve survival.

**Conclusion:** *PRMT5* KD epigenetically reduced TIC self-renewal, leading to increased survival in preclinical models. Pharmacological inhibition of PRMT5 enzymatic activity may have failed *in vivo* due to insufficient reduction of PRMT5 activity by chemical inhibition, or this failure may suggest that non-enzymatic activities of PRMT5 are more relevant.

**Implications:** Our findings show the importance of *PRMT5* to maintain and promote the growth of stemlike cells that initiate and drive tumorigenesis in pediatric high grade glioma.

## Introduction

As a group, PHGG is the deadliest childhood tumor.^1^ Five-year survival rates range from 2-20% depending on the subtype. PHGG is highly invasive and grows diffusely among normal cells, which limits surgery as a therapeutic option. Radiation therapy (RT) is transiently effective, but tumors almost always recur. Despite hundreds of clinical trials of cytotoxic and targeted chemotherapy, radiation remains the only accepted post-surgery standard-of-care therapy.

Epigenetic regulation plays an important role in PHGG tumorigenesis. The two diffuse PHGG subtypes are defined by epigenetic alterations. In diffuse midline glioma (DMG), histone 3 lysine 27 is altered to methionine (H3K27M); diffuse hemispheric glioma has a defining mutation of H3 glycine 34 to arginine or valine (H3G34R/V).^2,3^ These histone mutations prevent post-translational modifications, such as methylation, at the affected site. In DMG, for example, the H3K27M alteration prevents trimethylation of H3K27 (H3K27me3), resulting in the failure to silence expression of many genes and producing a broad group of oncogenic changes in transcriptional regulation.^4^ PHGG also includes a non-mutant histone 3 subtype that, despite being wild type for histone variants, may nonetheless rely on epigenetic regulation for oncogenesis and tumor growth.^5,6^

We conducted a short hairpin RNA (shRNA)-based screen of known epigenetic regulators in H3K27M mutant^7^ and non-mutant PHGG cell lines to identify genes that are important for tumor cell survival. The screening identified *PRMT5*, whose protein product methylates both NH_2_ groups of arginine’s guanidino functional group (known as symmetric dimethylation), as essential to PHGG cell growth. Methylation by PRMT5 occurs throughout the proteome and its targets include both histone and non-histone proteins.^8^ During normal mammalian development, PRMT5 maintains pluripotency of stem cell populations by upregulating expression of stemness genes and repressing genes associated with differentiation.^9^ Stem cell differentiation is accompanied by downregulation of PRMT5 expression.^9^ PRMT5 regulates transcription by methylating histone 3 arginine residues H3R2 and H3R8.^10,11^ This inhibits formation of H3K27me3, which produces a permissive environment for transcription of nearby genes.^10^ PRMT5’s arginine methyltransferase activity thus promotes expression of self-renewal genes otherwise silenced by H3K27me3 occupancy.^10^ PRMT5 also indirectly regulates the transcriptional activating mark H3K4me3. Here, PRMT5’s effect is context-dependent and can increase or decrease H3K4me3 levels.^12,13^

In cancers that depend on stem-like tumor cells for growth, PRMT5 has emerged as an important oncogene. PRMT5 expression correlates with increased tumor aggressiveness (grade) in adult high-grade glioma (AHGG) and promotes AHGG cell growth *in vitro.*^14^ PRMT5 also maintains self-renewal of tumor initiating cells and promotes tumor growth in preclinical models of breast cancer.^11^ In leukemia cell lines, PRMT5 promotes cell cycle progression through crosstalk-mediated inhibition of H3K27me3, which maintains the expression of genes that would otherwise be transcriptionally inactive.^10^

PHGG has a stem-like phenotype, suggesting that it originates from stem-like tumor-initiating cells (TICs).^5,15–21^ TICs are self-renewing and form progeny that divide rapidly and uncontrollably, forming highly invasive tumors.^15,20,22^ TICs are essential for tumor growth, resist current therapeutics, and may be a primary cause of treatment failure.^16,20,23–25^ We hypothesized that PRMT5 promotes PHGG oncogenesis by promoting TIC formation and growth.

We investigated PRMT5’s role in PHGG, both *in vitro* and *in vivo.* PRMT5 regulates cell self-renewal *in vitro* and its genetic depletion suppresses PHGG xenograft growth *in vivo.* ChIP-Seq analysis showed PRMT5 mediates epigenetic alterations to H3K27me3, H3K27ac, and H3K4me3 occupancy at genes important to maintain stemlike characteristics.

## Methods

### Cell culture

PHGG cell lines BT-245, SU-DIPG-IV (DIPG4), HSJD-DIPG-7 (DIPG7), SU-DIPG-XIII (DIPG13), and HSJD-GBM-001 (GBM1) were cultured in suspension in low-attachment 250 mL/75 cm^2^ flasks (Greiner #658, Supp. Table 1). The medium for DIPG4, DIPG7, DIPG13 and GBM1 consisted of equal parts of DMEM/F12 (Gibco) and Neurobasal-A (Gibco), 2% of B-27 growth supplement without vitamin A (Gibco), 1% each of L-Glutamine (Gibco), HEPES (GIbco), non-essential amino acids (Gibco) and sodium pyruvate (Gibco) and antibiotic/antimycotic (Gibco), 0.1% heparin (StemCell Technologies), and EGF, FGF and PDGF A/B at 20 ng/μL (Shenandoah Biotechnology). The medium for BT-245 consisted of Neuro-cult with proliferation supplement (StemCell Technologies), 1% of pen-strep antibiotic (GIbco), 0.1% of heparin (StemCell Technologies), and of EGF, FGF and PDGF A/B at 20 ng/μL (Shenandoah Biotechnology). Cells were passaged as required by trituration to break up the neurospheres and replacement of the medium.

### PRMT5 *KD*

We obtained three shRNAs targeting *PRMT5* in a lentiviral format (TRCN0000379612, TRCN0000303446, and TRCN0000381130) and an empty vector control (SHC202) from the Functional Genomics Facility at the University of Colorado Anschutz Medical Campus. Cells to be transduced were placed in 24-well ultra-low attachment plates (Corning #3473) at 100,000 cells in 2 mL of medium. Polybrene (1 µg/mL) was added to the cell suspension and, 20-30 minutes later, medium was added containing lentiviral shRNA particles at an MOI of 0.25-0.33. Following 4-6 hours of incubation, the medium was changed and the cells were incubated for 48-72 hours. After 72 h, puromycin (1 µg/mL) was added. Cells were selected for one or more passages prior to beginning the experiments.

### shRNA screen

We conducted an shRNA screen in two cell lines, SU-DIPG-IV, derived post-autopsy from a pontine tumor with H3.1K27M and ACVR R328V mutations (DIPG4)^7^, and VUMC-DIPG-10, derived post-autopsy from an H3K27-wild type pontine tumor with an NF1 Q209* variant and *MYCN* amplification (DIPG10). The shRNA screening library comprised 4,139 shRNAs targeting 405 genes involved in the epigenetic regulation of gene expression (Supp. Data 1-2). Cells were transduced with the barcoded shRNA library at an MOI calculated to obtain one shRNA per cell and placed into selection. Samples were collected after 72 hours of selection and 21 days later. The barcodes were amplified and sequenced, and the resulting data were analyzed to identify changes in barcode frequency in the samples.

### Cell growth assay

Cells previously transduced with the empty vector control or shRNA targeting *PRMT5* were triturated to obtain a single-cell suspension and placed into 24-well plates (100,000 cells per well). In long-term longitudinal assays, cells were triturated into single cells and counted once or twice per week for up to five weeks. Following each counting, 100,000 cells were replated. In short-term longitudinal assays, 10,000 cells transduced with empty vector control or one of three *PRMT5* KD constructs were placed into 96-well plates in triplicate. Cells were then incubated in an Incucyte S3 Live Cell Analysis Instrument (Sartorius) for six days, during which imaging was conducted every four hours. Image data were analyzed to ascertain cell growth based on a neurosphere proliferation model (Sartorius).

### Extreme-limiting dilution assay of self-renewal

Cells were triturated to form a single-cell suspension and placed into 100 μL of medium in a 96-well plate at 1 cell per well (16 wells), 2 cells per well (8 wells), 4 cells per well (8 wells), 8 cells per well (8 wells), 16 cells per well (8 wells), 32 cells per well (8 wells) or 64 cells per well (8 wells). Cells were incubated for 2-3 weeks, and wells with neurospheres were identified using a brightfield microscope that accepts a 96-well plate (Keyence). Results were analyzed as previously described to yield point estimates of mean stem cell frequency and a p-value comparing *PRMT5* KD to empty vector control cells.^26^

### Apoptosis assay

Cells were placed into 96-well white wall plates at 2,000 cells per well with three technical replicates per sample. Caspase-Glo 3/7 3D Assay (Promega, #G8981) was added and the mixture incubated per manufacturer instructions. After incubation, luminescence was measured using a Synergy Plate Reader (BioTek).

### Cell cycle analysis

Cells were fixed in 70% ethanol at 4C for at least 12 hours. Cells to be analyzed were placed into a 96-well plate at 100,000 cells per well in 200 µL of medium, spun down, rinsed with PBS, and resuspended in Guava Cell Cycle Staining Reagent (Millipore). Data acquisition was performed on a Guava PCA-96 system (Millipore). Data were analyzed using FlowJo (FlowJo, LLC) to identify cells in G1, S, and G2 cell cycle phases.

### Bulk RNA-Seq

We conducted bulk RNA-Seq of *PRMT5 KD* and empty vector control cells. Samples were obtained from BT245, DIPG4, DIPG7, DIPG13, and GBM1 cells and comprised one empty vector control and three KD samples with different shRNA vectors. RNA extracted from 1×10^6^ cells was sent to the Anschutz Genomics Shared Resource for sequencing. Sequencing libraries of polyA RNA were prepared using the TruSeq Library Preparation Kit v2 (Agilent). Paired-end sequencing, with 50 million reads per sample, was performed using a NovaSeq 6000 (Illumina). Data were mapped to the human genome (GRCh38) using gSNAP. Expression (FPKM) was derived using Cufflinks, and differential expression was analyzed with ANOVA in R. FPKM read data were analyzed using gene set enrichment analysis (GSEA) and Metascape.

### ChIP-Seq

ChIP-Seq was performed using antibodies against H3 lysine 4 trimethyl (H3K4me3, Cell Signaling Technology (CST) #9751S, 1:50), H3 lysine 27 trimethyl (H3K27me3, CST #9733S, 1:50), and H3 lysine 27 acetyl (H3K27ac, CST #8173S, 1:50). We also obtained and sequenced an input sample and a sample immunoprecipitated with Rabbit IGG only (CST #3900S, 2.5 ug total antibody per reaction). Approximately 5-10 million cells were fixed in 1% formaldehyde, after which cells were lysed using FL lysis buffer.^27^ Nuclei were sonicated at low energy to fully isolate them from the cytosol. Nuclei were then sonicated at high energy in a Bioruptor Plus (Diagenode). Sonication parameters to produce DNA fragments in the 200-600 bp range were optimized beforehand. Samples were incubated with antibody overnight at 4C with rocking. Magnetic beads (CST #9006) were used for immunoprecipitation. DNA was eluted from the beads using 2X ChIP Elution Buffer (CST #7009) and incubating at 65C for 30 min with shaking. Following elution, DNA was purified using spin columns (CST #14209). DNA was sequenced on a NovaSeq 6000 (Illumina) using 150 paired-end cycle reads with 50 million paired reads per sample. Following filtering, reads were aligned to GRCh38 using Bowtie. Bams were filtered to remove secondary alignments, improperly paired reads, and alignments with mapping quality <30. Peak calling was performed using MACS3.^28^ Peaks were analyzed using DiffBind^29^ (3.8.4) and ChIPpeakAnno^30^ (3.32.0) in R (4.2.2), followed by differential gene expression, GSEA and Metascape analyses.

### Tumor implantation in mice

All animal experiments were conducted in accordance with an existing IACUC protocol administered by the University of Colorado Anschutz Medical Campus. BT-245 (a DMG cell model) or GBM1 (a cortical GBM model) cells genetically modified to express a luciferase (Luc) gene were implanted into the brains (right pons for BT-245 and right striatum for GBM1) of athymic nude mice (Charles River) by intracranial injection using a stereotactic frame.^31^ Following injection, the mice were monitored daily for evidence of illness, including decreased activity, hunched posture, poor grooming, and ataxia. In addition, mice were weighed weekly. Animals that were moribund and/or had lost more than 15% of their initial body weight were euthanized for necropsy. Tumor growth was monitored by MRI, as well as bioluminescence imaging (BLI) for cell lines recombinantly modified by the introduction of a Luc gene. In experiments involving the injection of *PRMT5* KD cells, differences in survival were assessed. In experiments involving radiation therapy (RT) and drug treatment with PRMT5 inhibitor LLY-283 (SelleckChem), survival was evaluated in four experimental arms: vehicle control, RT only, LLY-283 only and LLY-283 plus RT. We used eight mice per arm in each experiment, which enabled the detection of a 50% survival difference at 80% power with an alpha of 0.05.

### In vitro cell survival experiments

Cells were placed into 96-well plates at 20,000 cells per well in 90-100 μL medium with three replicates per condition. In experiments utilizing radiation therapy (RT), cells were irradiated using a Cs source. In experiments using the PRMT5 inhibitor LLY-283, drug dissolved in DMSO and diluted in phosphate buffer solution was added to each well at concentrations ranging from 0.316 nM to 10 μM by half-log_10_ concentration increments. Cells were then incubated for 120 hours. Following incubation, CellTiter 96 Aqueous One Solution Assay (Promega) was added (20 μL per well), and cells were incubated 1-4 hours. Plates were then analyzed for absorbance at 490 nM in a 96-well Synergy plate reader (Biotek).

### Data availability

Sequencing data (bulk RNA-Seq and ChIP-Seq) are being deposited in the NCBI Gene Expression Omnibus (GEO) database. [Assignment of an accession (GSE) number is pending and will be updated when the submission is complete].

## Results

### *PRMT5* is essential for PHGG cell growth

An shRNA KD screen identified epigenetic regulators that are important for tumor cell growth in DMG^7^ and H3K27 wild-type (wt) pontine PHGG cell lines (Figure 1a, Supp. Figure 1a). Reanalysis of the DMG screen in conjunction with the previously unpublished H3K27 wt PHGG screen revealed two epigenetic regulators, *PRMT5* and *HDAC2*, with significant negative fold change in both cell lines (Figure 1a, Supp. Table 2). Based on DEMETER2 RNA interference (RNAi) screening data (Broad Institute) in normal and non-pediatric cancer cell lines, *PRMT5* was the more attractive target. The DEMETER2 data show *PRMT5* promotes growth of adult glioma and other central nervous system (CNS) tumors (Figure 1b), while *HDAC2* promotes tumor growth in the peripheral nervous system but not the CNS. To exclude the possibility that the *PRMT5*-related growth effect occurred because *PRMT5* is essential for survival of all cells, we compared DEPMAP CRISPR knockout (KO) data (Broad Institute) with DEMETER2 RNAi knockdown (KD) screening results of *PRMT5* (Figure 1c) in cancer and non-cancer cell types.^32^ In KO experiments, cells without *PRMT5* failed to survive; however, in KD experiments, normal cells survived *PRMT5* KD, but cancer cells reliant on *PRMT5* expression did not survive. Accordingly, while a baseline level of *PRMT5* expression is essential for the survival of all cells, the expression level of *PRMT5* in KD experiments was sufficient to enable the survival of non-cancer cells. We verified that targeting *PRMT5* with shRNA reduced, but did not eliminate, its gene and protein expression in PHGG cells (Figure 1d-e, Supp. Figure 1b). A functional assay showed decreased symmetric dimethylation of arginine residues (PRMT5’s sole enzymatic activity and target) as a result of *PRMT5* KD (Figure 1f). Symmetric dimethylation also decreased when PHGG cells were treated with the clinical-grade PRMT5 inhibitor LLY-283 (Figure 1g).

**Figure 1.**
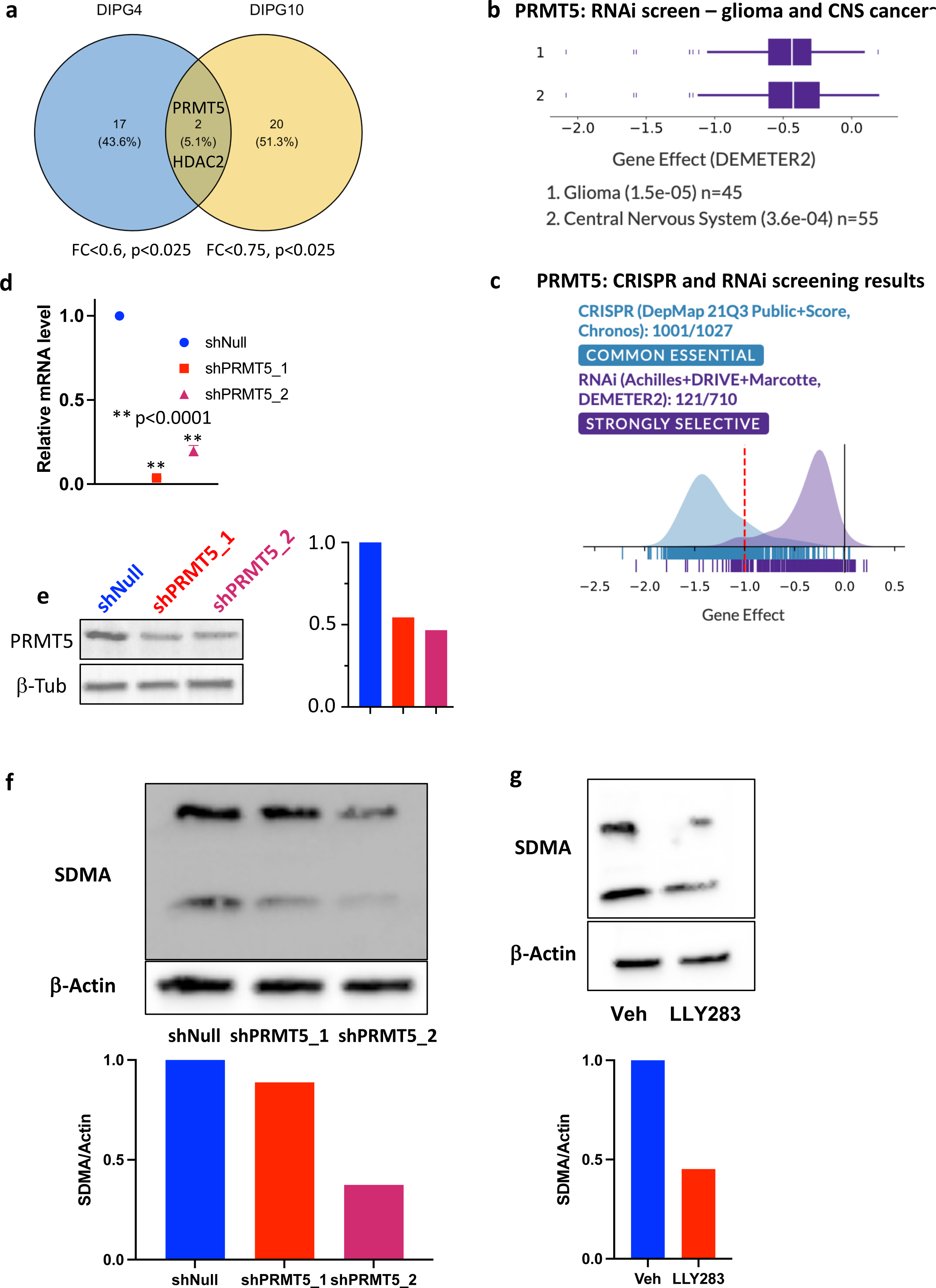
PRMT5 is essential for growth of PHGG and other CNS cancer cells. a. shRNA KD screen of epigenetic regulators identified those that are important for cell growth; Venn diagram shows genes important for tumor cell survival in DIPG4 (H3K27M DMG cell line), DIPG10 (H3-wt pontine PHGG) and overlap between the two cell types; b. In RNA interference (RNAi) knockdown (KD) screens, CNS cancer and glioma cell lines depend on *PRMT5* to proliferate, validating PRMT5 as a potential therapeutic target in these cancer types; c. DepMap CRISPR knockout (KO) screening shows PRMT5 is essential for cell survival at a level similar to housekeeping genes in 97.5% (1001/1027) of cell lines tested, suggesting PRMT5 KO is lethal to cancer and non-cancerous cell types; in RNA interference knockdown (KD) screening, however, *PRMT5* KD is “strongly selective” among different cell types, suggesting it may slow growth of cancer cells without adversely affecting normal cells; d. Lentiviral transduction of cells with shRNA targeting *PRMT5* produces KD levels of 80-90%; e. Western blot (left panel) showing relative PRMT5 protein levels following shRNA mediated KD, Western blot quantification (right panel) shows *PRMT5* KD produces protein expression decrease of 45-55%; f. Western blot (upper panel) of symmetric dimethylation of arginine residues throughout the proteome shows the effect of *PRMT5* KD on PRMT5 enzymatic activity; Western blot quantification (lower panel) shows relative levels of SDMA expression following *PRMT5* KD; g. Western blot (upper panel) shows SDMA levels following treatment of GBM1 cells with the clinical-grade PRMT5 inhibitor LLY283; Western blot quantification (lower panel) shows effect of LLY283-mediated inhibition of SDMA levels.

### *PRMT5* KD reduces self-renewal of PHGG stem-like cells, reduces proliferation and cell cycle progression, and increases apoptosis

We evaluated the effects of *PRMT5* KD and PRMT5 inhibition using LLY283 on PHGG cells. In *in vitro* extreme limiting dilution assays (ELDA), *PRMT5* KD and LLY283 both decreased the tendency of PHGG cells to form neurospheres. We calculated that *PRMT5* KD/inhibition decreased the frequency of self-renewing cells by a factor of 6.6 or more (p<7e-9, Figure 2a-b). We interrogated a single-cell RNA-Seq dataset of PHGG patient samples and identified a likely TIC population that expressed *NES, SOX2* and *OLIG2,* which are also expressed in neural stem cells or oligodendrocyte progenitors (Supp. Figure 1c). These undifferentiated cells suggest a likely target for PRMT5 activity. *PRMT5* KD decreased PHGG cell growth in both long-term and short-term cell growth assays (Figure 2c, Supp. Figure 1d). The effect of PRMT5 inhibition using LLY283 on PHGG cells was dose-dependent, with a significant portion of cells (approximately 75%) remaining viable even at the highest dosage levels (Figure 2d). LLY283 showed no dose-dependent effect on survival when applied to a normal human astrocyte cell line (Supp. Figure 1e). *PRMT5 KD* inhibited cell cycle progression, increasing the number of cells in the G1 phase and decreasing the number of cells in the S or G2 phases (Figure 2e). *PRMT5 KD* also consistently increased apoptosis by 4-5% in *in vitro* experiments (Figure 2f).

**Figure 2.**
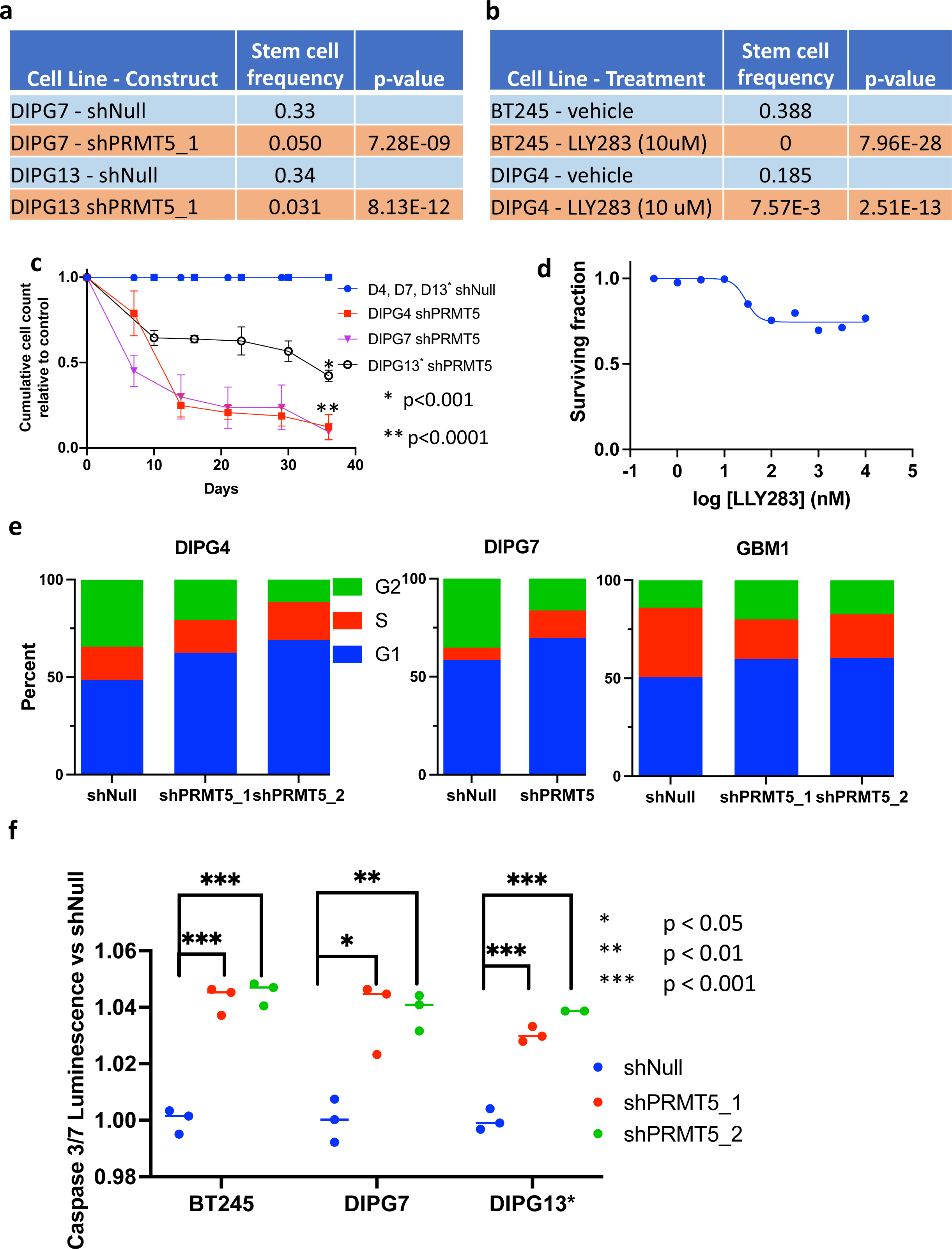
*PRMT5* KD decreases self-renewal, inhibits cell growth and cell cycle progression, and increases apoptosis versus empty vector control. a. In extreme limiting dilution assay, *PRMT5* KD decreases stem cell frequency (measure of self-renewal) in PHGG cell lines; b. PRMT5 inhibition using LLY283 decreases stem cell frequency in PHGG cell lines; c. *PRMT5* KD results in decreased cell growth in DMG cell lines; d. PRMT5 inhibition using LLY283 has a dose-dependent effect on survival but leaves a large resistant population; e. *PRMT5* KD slows cell cycle progression by increasing cells in G1 phase and decreasing cells in G2 phase; f. *PRMT5* KD increases cell susceptibility to apoptosis.

The overall effects of PRMT5 KD/inhibition *in vitro* thus included decreased PHGG TIC frequency, stem cell gene expression, and cell growth. *PRMT5* KD also inhibited PHGG cell cycle progression and increased apoptosis. These effects are consistent with the hypothesis that *PRMT5* maintains TIC self-renewal in PHGG.

### *PRMT5* KD increased expression of genes associated with stem cell maintenance and oncogenesis

We conducted bulk RNA-Seq on *PRMT5 KD* versus control cells. Data were analyzed by taking the two KD samples with the lowest expression levels of *PRMT5* and comparing them to the empty vector control for each cell line. Differentially expressed genes as a result of *PRMT5* KD included several genes involved in stem cell maintenance (*PAX3, CDH4, KIF1A, and UCN2*) as well as oncogenesis (*CDH4, MMP14, and ARPC1B*) (Figure 3a, Supp. Table 3). We conducted pathway analyses using geneset enrichment analysis (GSEA) and Metascape.^33–35^ GSEA demonstrated that *PRMT5* KD samples were depleted in gene sets of core stem cell genes and enriched in gene sets expressed in differentiating cells (Figure 3b). Our GSEA results also showed that *PRMT5* KD cells were enriched in DNA repair pathways and depleted in genes expressed during hypoxia and in mesenchymal cells (Figure 3b). Metascape analysis was consistent with the GSEA results (Supp. Figure 2a-b).

**Figure 3.**
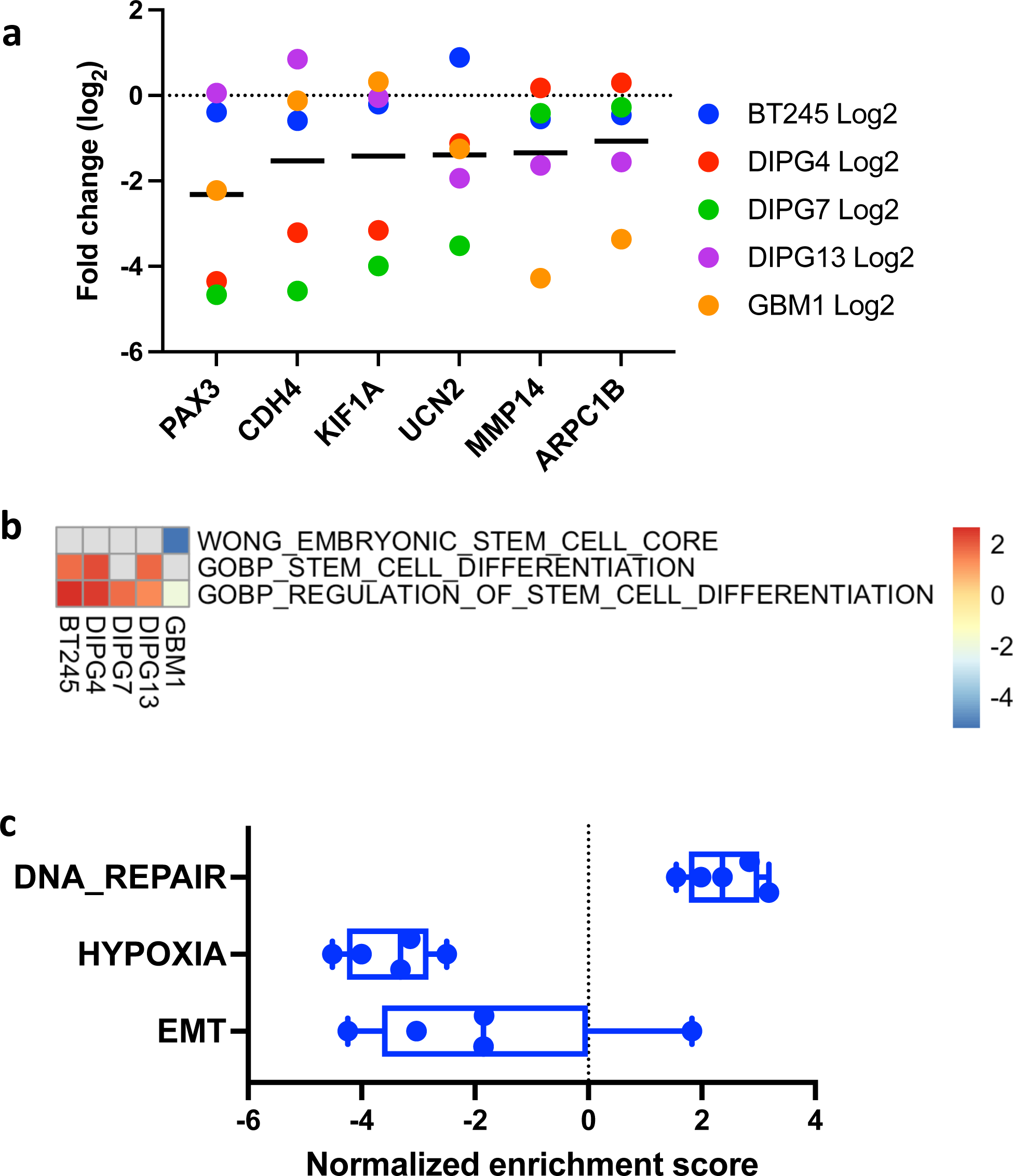
*PRMT5* KD decreases expression of genes important to stem cell growth and decreases expression in important cancer pathways. a. Log2 fold change following *PRMT5* KD in five cell lines for six representative stemness/cancer genes; black bar represents mean value; b. Heatmap of GSEA results showing Net Enrichment Score (NES) for stemness-related gene sets for *PRMT5* KD versus empty vector control in PHGG cell lines; gene ranking for GSEA based on log2 value of ratio of average *PRMT5* KD gene expression divided by empty vector control value (False discovery rate (FDR) <0.25 for all NES values); c. GSEA-based NES values for gene sets important for PHGG inception and growth (FDR <0.05 for all NES values).

Similar to its phenotypic effects, *PRMT5* KD decreased the expression of genes important for stem cell maintenance and oncogenesis in PHGG cells. On a pathway level, *PRMT5* KD decreased self-renewal, increased differentiation and altered oncogenic pathways in a direction of lessened tumor growth.

### *In vivo* studies of *PRMT5 KD* in PHGG showed increased survival and decreased tumor aggressiveness

We orthotopically implanted *PRMT5* KD cells and empty vector control cells (BT245) into mice and conducted a survival study. *PRMT5* KD mice survived significantly longer than control mice; however, all mice eventually died of tumor-related effects (Figure 4a). MRI and bioluminescence imaging confirmed that the *PRMT5* KD tumors were less aggressive than the control tumors (Figure 4b-c). Hematoxylin and eosin (H&E) staining showed that the tumor extent and cell density were lower in *PRMT5* KD cells (Figure 4d). Staining for Ki-67 verified that the frequency of proliferating cells was lower in *PRMT5* KD cells than in control tumor cells (Figure 4e).

**Figure 4.**
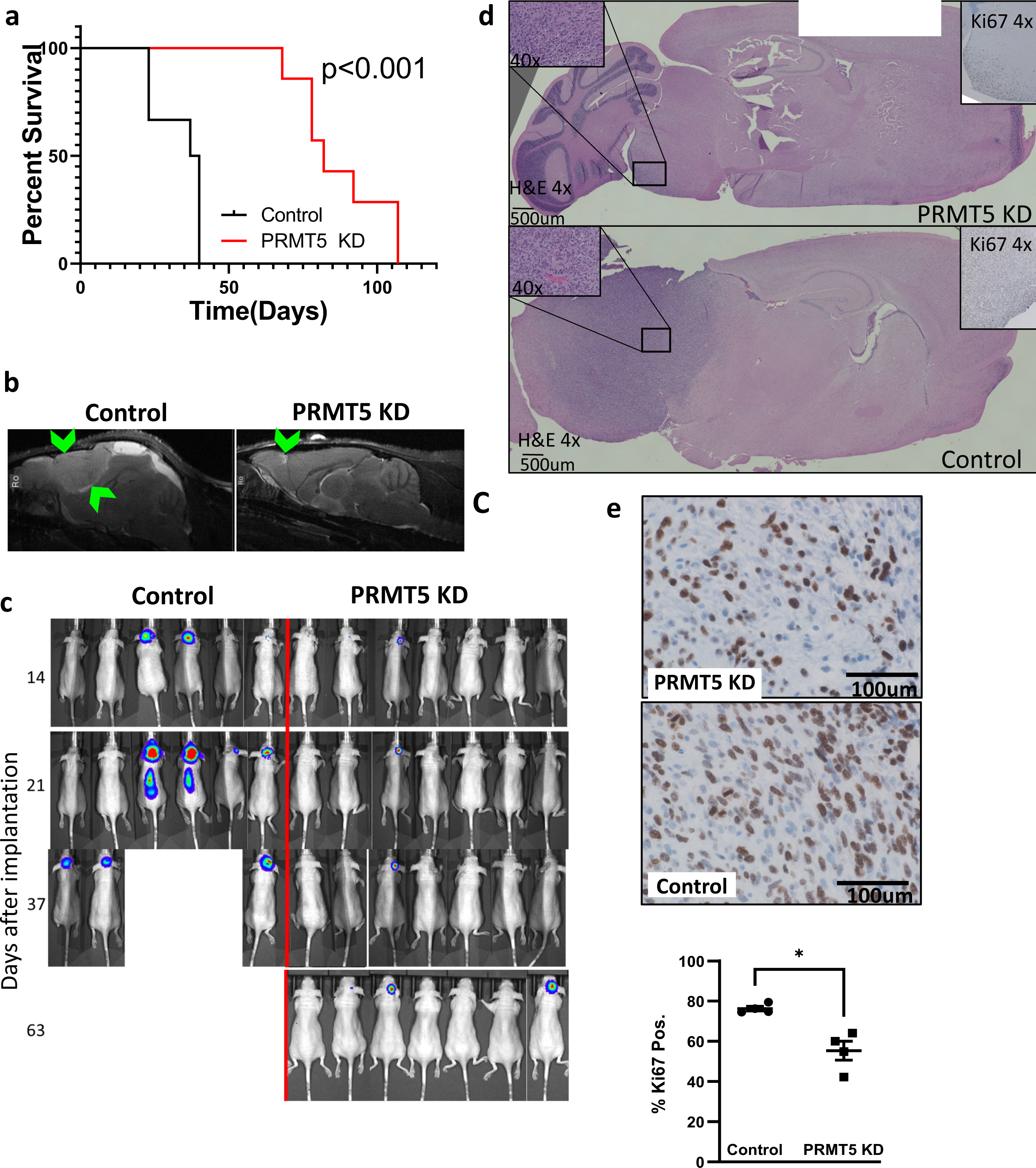
*PRMT5* KD produces less aggressive tumors with greater survival in mouse PDX model of PHGG. a. Kaplan-Meier survival curve showing that mice injected with tumor cells with *PRMT5* KD survive 2.5-3 times longer than mice injected with empty vector control tumor cells; b. MRI images of empty vector (control) and *PRMT5* KD tumors; arrowheads show enhancement of tumor areas; c. Bioluminescence images showing tumor progression; d. Histological sections with H&E staining showing representative tumors from *PRMT5* KD and empty vector control mice; e. Ki-67 staining (upper panels) and quantification, showing greater number of proliferating cells in *PRMT5* KD versus empty vector control tumors (p<0.05).

We also conducted a mouse survival study using PRMT5 inhibitor LLY283 and the current standard of care therapy, RT. *In vitro* experiments combining RT with *PRMT5* KD showed 17.1-30.3% lower survival rates when *PRMT5 KD* versus control cells were exposed to RT (Figure 5a-b). These results suggested that *PRMT5* depletion sensitizes cells to the effects of RT. Because a large fraction of tumor cells survived *in vitro* treatment with LLY-283 alone (Figure 2d), we hypothesized that combining PRMT5 inhibition with RT might lead to increased cell death, but that LLY283 alone would be ineffective. Accordingly, we designed a mouse experiment to apply RT when BLI showed a signal indicating early-stage tumor engraftment. We found no significant differences in survival among any of the four treatment groups (Figure 5c-d).

**Figure 5.**
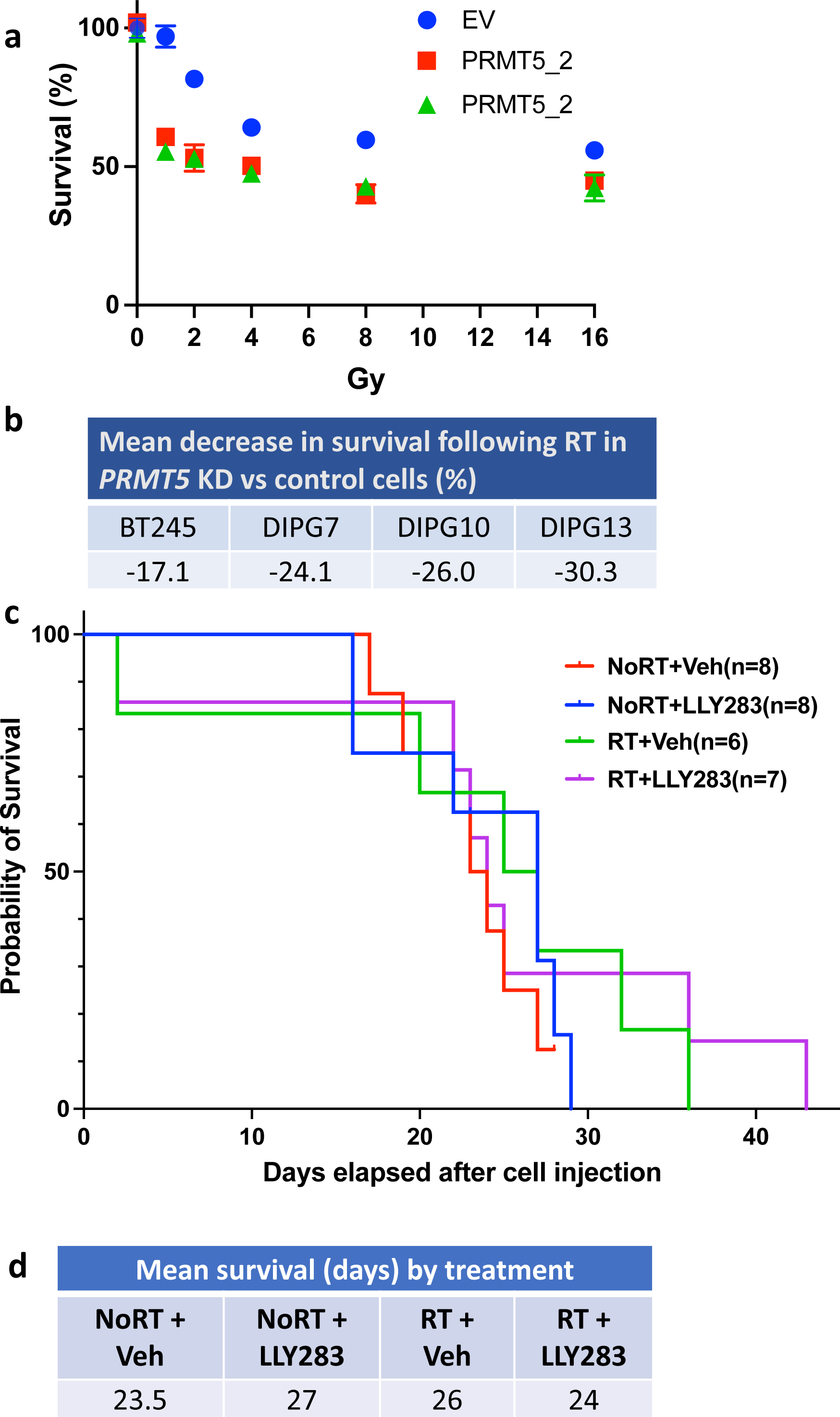
*PRMT5* inhibition did not result in greater survival in mouse PDX model of PHGG. a. In *vitro*, PHGG cells with *PRMT5* KD are more susceptible to RT than empty vector control cells; b. Quantification of decreased survival (%) of *PRMT5* KD versus empty vector control cells from panel a; c. Kaplan-Meier survival curve showing no significant differences in survival of mice treated with PRMT5 inhibitor versus vehicle control with and without RT.

*PRMT5* KD in our mouse PDX experiment had effects similar to those observed *in vitro*. However, the combination of RT and *PRMT5* inhibition did not increase survival. As further explored in the Discussion, the reasons for treatment failure could be attributed to the beginning of inhibitor treatment too late, that is, after the tumor had already passed through the stage where a stemlike population is required for tumorigenesis, or could have resulted from other factors.

### *PRMT5* KD alters chromatin occupancy at important epigenetic regulatory sites

We performed ChIP-Seq in the two cell lines used in the mouse PDX experiments (BT245 and GBM1). We immunoprecipitated H3K4me3, H3K27ac, and H3K27me3, performed high-throughput sequencing of the extracted DNA, and analyzed the changes in chromatin occupancy (Figure 6, Supp. Figures 3-7). Overall, occupancy at the H3K27me3 and H3K4me3 marks decreased, while occupancy at the H3K27ac mark decreased only slightly with *PRMT5* KD (Figure 6a-b). Principal component analysis comparing occupancy changes showed that tumor type (H3K27-mutant (BT245) versus H3K27-wt (GBM1)) explained the greatest amount of occupancy differences at all three histone marks (PC1, 60-91%), whereas *PRMT5* KD was responsible for the next greatest amount of occupancy variation (PC2, 4-15%) (Supp. Figure 3g).

**Figure 6.**
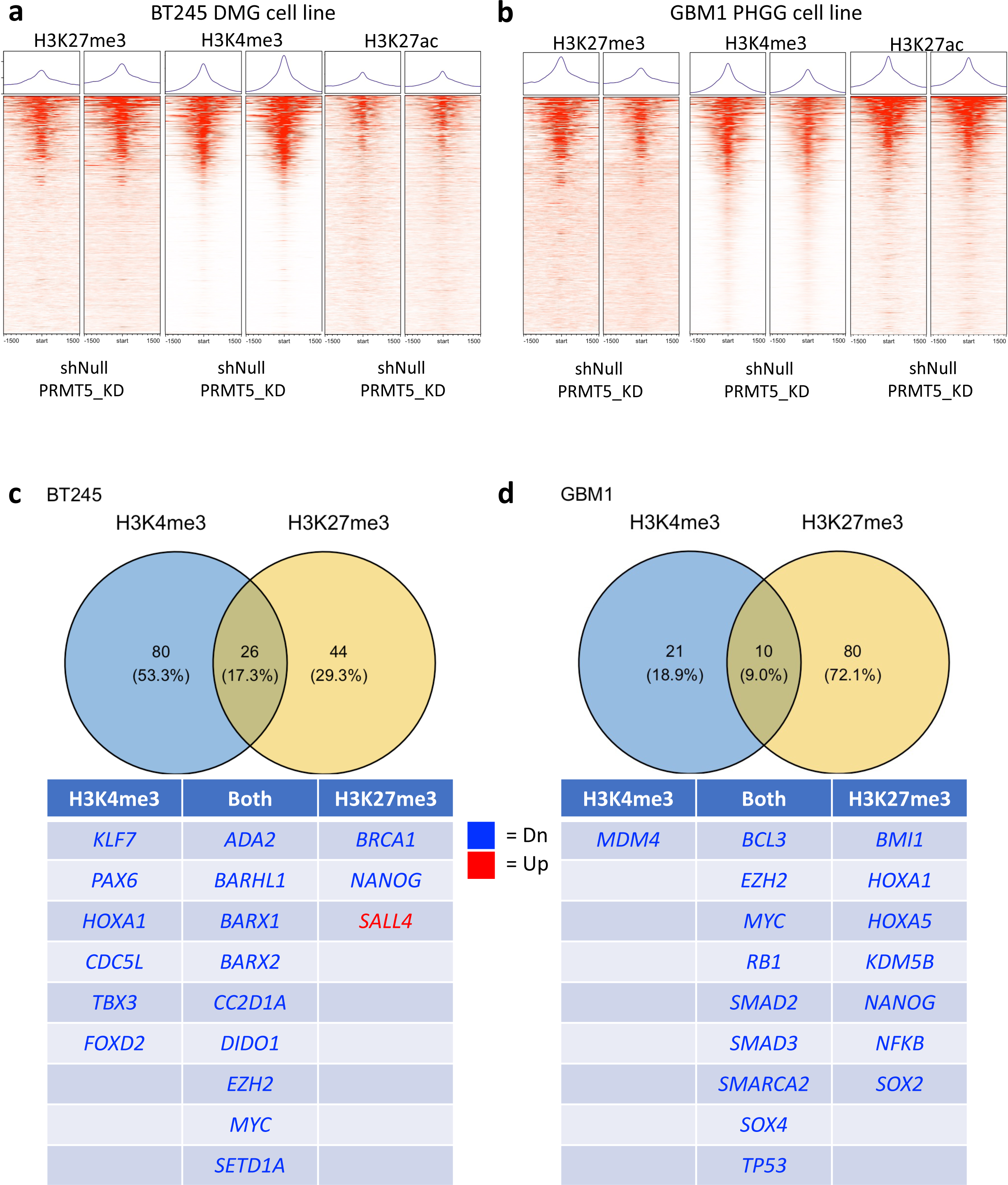
ChIP-Seq in *PRMT5* KD versus empty vector control shows consistent differences in H3K27me3 and H3K4me3 occupancy. a. – b. occupancy levels of 1,000 randomly selected genes from 1,500 bp downstream (-1,500) to 1,500 bp upstream of transcription start site (TSS) in *PRMT5* KD and empty vector control cells; panel a shows BT245 cells; panel b shows GBM1 cells; c. – d. Upper panels: Venn diagram of GSEA results showing number of target gene sets with significant occupancy changes as the result of *PRMT5* KD at H3K4me3 (left) or H3K27me3 (right) histone marks, overlap shows gene sets with significant occupancy changes at both marks; lower panels: GSEA results with target gene sets listed by transcription factor active in stem cell maintenance or differentiation corresponding to Venn diagram regions, panel c – BT245 cells; panel d – GBM1 cells, blue type = depletion; red type = enrichment.

Next, we identified binary changes in occupancy (occupied/unoccupied) by genomic locus for the three samples (two *PRMT5* KD constructs versus one empty vector control) used in each immunoprecipitation reaction (Supp. Figure 3a-f, 4a-f). To understand the transcriptional effects of *PRMT5* KD, we performed GSEA on the ChIP-Seq results using the hallmark, curated and ontology gene sets provided in the MSigdb, ranking genes by change in histone mark peak height attributable to *PRMT5* KD.^33,34,36^ We identified gene sets with “target” in their names because they typically represent a set of target genes acted upon by a single transcription factor (TF) (Figure 6c-d; Supp. Tables 4-7). In both cell lines, occupancy at these “target” sites overwhelmingly decreased at both the H3K4me3 (BT245: 106/106 sites with FDR<0.05; GBM1: 30/31 gene sets with FDR<0.05) and H3K27me3 marks (BT245: 62/70 gene sets with FDR<0.05, GBM1: 90/90 gene sets with FDR<0.05) (Figure 6c-d, Supp. Tables 4-7). Target gene sets representing TFs involved in early embryonic development or stemness maintenance in highly undifferentiated cells decreased in peak height at H3K4me3 with *PRMT5* KD, while decreases in H3K27me3 occupancy tended to occur at targets of TFs associated with more differentiated cells (Figure 6c-d). A few genes important for stem cell maintenance, for example *SOX2* in GBM1 cells, decreased in H3K27me3 occupancy as the result of *PRMT5* KD (Figure 6d). This decrease does not necessarily equate to increased transcription, but instead may mark a change in the *SOX2* locus to being poised for transcriptional silencing as differentiation occurs.^37^ 17.3% of the targets in BT245 and 10% of targets in GBM1 decreased at both H3K4me3 and H3K27me3 (Figure 6dc-d). Bivalent shifts at these loci may likewise suggest a shift from higher transcriptional levels to a loci poised for silencing upon cellular differentiationr.^37,38^

Differential gene occupancy analysis identified several consistent enrichment or depletion differences (Supp. Figure 5a). These included genes of the *SEMA4* family that are involved in cellular differentiation and have been identified as important to the development of cancer^39^; *CACUL1*, which regulates the G1/S cell cycle transition^40^; *PWWP2A*, which recruits histone deacetylases to gene promoters and is essential for mitosis^41^; *SMURF1*, a ubiquitin ligase specific to bone morphogenetic proteins^42,43^; and *TOP3B*, a topoisomerase involved in recombination that has oncogenic characteristics^44^ (Supp. Figure 5a). Pathways suggested by these occupancy differences included: regulation of cellular proteins through destruction (ubiquitination and the clathrin pathway), bone morphogenetic protein regulation, cell division, and transcriptional regulation (DNA repair and RNA polymerase II regulation), cellular processes important for oncogenesis such as differentiation and angiogenesis, and the cellular stress response (heat shock proteins) (Supp. Figure 5b). We performed ontological analysis using Metascape on the ChIP-Seq data for each of the two cell lines (Supp. Figures 6-7). The gene ontology (GO) terms from the Metascape analysis, which included cellular signaling, ligand/receptor interaction, differentiation and cell fate commitment, angiogenic signaling, and brain development, were consistent with the GSEA and differential expression analyses (Figure 6c-d, Supp. Figures 6-7).

Analysis of the ChIP-Seq data suggests that transcriptional regulation through the H3K27me3 and H3K4me3 histone marks is an important aspect of PRMT5’s maintenance of stemlike TIC cells important for the development of PHGG. Genes whose occupancy decreased at H3K4me3 with *PRMT5* KD were primarily those involved in stem cell maintenance. Occupancy changes at H3K27me3 following *PRMT5* KD were consistent with the inception of differentiation. The differential gene expression and Metascape analyses were consistent with the GSEA results in showing that *PRMT5* regulates stem cell characteristics in DMG and PHGG through modulation of the H3K4me3 and H3K27me3 chromatin marks.

## Discussion

Our results show that PRMT5 plays an important role in the development and growth of PHGG through epigenetic control of gene expression in stem-like TICs. PRMT5’s regulation of TIC self-renewal and growth underlies the phenotypic effects of *PRMT5* KD in PHGG. Those phenotypic effects include decreased proliferation and cell cycle progression, increased susceptibility to apoptosis, and decreased self-renewal. Genes whose expression *PRMT5* KD affected include *PAX3*, whose expression markedly diminished. *PAX3* plays key roles in neural stem cell maintenance, neurogenesis, and astrogenesis.^45,46^ GSEA of the RNA-Seq data showed that *PRMT5* KD downregulated pathways that maintain self-renewal and other stem cell characteristics and increased expression in cellular differentiation pathways. *PRMT5* KD also decreased hypoxia and mesenchymal gene expression and increased DNA repair. These changes are associated with decreased tumorigenesis in DMG and H3K27-wt PHGG.

The phenotypic and gene expression changes that *PRMT5* KD induced occurred in both DMG cells bearing the H3K27M mutation and H3K27-wt PHGG. Our ChIP-Seq studies, which focused on alterations in trimethylation and acetylation of H3K27 and trimethylation of H3K4, offer some clues regarding the mechanism by which *PRMT5* KD inhibits PHGG growth in both H3K27-wt and H3K27M cells. The H3K4 and H3K27 methylation sites were relevant because *PRMT5,* though it encodes an arginine methyltransferase, regulates methylation through a cross talk-mediated mechanism.^10^ H3K4me3 occupancy decreased with *PRMT5* KD in genes that maintain self-renewal and other stem cell characteristics important to oncogenesis. In both H3K27-wt and H3K27M cells, *PRMT5* KD also produced changes in H3K27me3 occupancy that are consistent with cellular differentiation. An important question for DMG (H3K27M) cells is the extent to which changes in H3K27me3 are important to overall gene expression. Because the H3K27M mutation is estimated to be less than 20% penetrant in DMG, most H3K27 loci in DMG cells are non-variant.^4^ The H3K27M variant, however, sequesters the PRC2 complex that is responsible for adding the trimethyl mark at the H3K27 locus, decreasing the number of H3K27me3 sites to a greater extent than the penetrance of the H3K27M mark alone would suggest.^4^ The implications of the DMG ChIP-Seq results are that, first, the H3K27M mutation does not prevent PRMT5 from regulating H3K27me3 in DMG cells. The observation that *PRMT5* KD downregulated self-renewal and other stem cell characteristics and led to differentiation suggests that changes at the H3K27me3 affected transcription in H3K27M mutant cells.

At least some changes in occupancy at the H3K27ac mark also occurred in both cell lines. These changes involved some of the same genes identified in the H3K4me3 and H3K27me3 ChIP-Seq experiments, probably for two reasons. First, H3K27me3 and H3K27ac marks are mutually exclusive, meaning that one has to be removed for the other to occur, so changes in both are required in many cases. Second, the H3K27ac mark is associated with transcriptional enhancers and may, therefore, indicate increases or decreases in transcription, as opposed to the occurrence of conditions that favor or disfavor transcription. The fact that several genes identified as potentially important for PHGG formation and growth had differences in H3K27ac occupancy with *PRMT5* KD suggests that further investigation of the role of H3K27ac may be worthwhile.

Our ChIP-Seq results included H3K27me3 and H3K4me3 occupancy changes with *PRMT5* KD in ubiquitination, the bone morphogenetic protein (BMP) pathway, and the clathrin pathway. To date, these pathways have not received much attention in PHGG. Ubiquitination controls the degradation of intracellular proteins, which play key roles in protein regulation. Clathrin-mediated vesicular transport enables intercellular communication as well as intracellular protein transport. BMP signaling regulates early CNS development and patterning and, later, regulates fate specification of CNS cells, including the oligodendroglial-to-neuronal shift.^47^ BMP signaling is also involved in the shift from neuronal to astrocyte fate towards the end of neurogenesis.^47^ Platelet-derived growth factor (PDGF) signaling, which is an important driver of PHGG growth, is active in these same processes. The epigenetic changes introduced by *PRMT5* KD thus highlight previously underappreciated pathways that might be important and deserve further study in PHGG.

While our *in vivo* experiments showed a survival benefit and decreased aggressiveness with *PRMT5* as compared to empty vector control cells, PRMT inhibition with a clinical-grade drug showed no survival or phenotypic differences in mice. This was true whether the inhibitor was administered alone or in conjunction with RT. Another group also recently reported that PRMT5 inhibition, given without RT, failed to confer a survival benefit.^48^ One possible reason the inhibitor failed could be that *PRMT5* is not acting enzymatically but is instead playing a non-enzymatic role such as acting as a scaffold protein. Inhibition of PRMT5’*s* enzymatic activity would not be expected to produce a therapeutic effect if PRMT5 were acting in a non-enzymatic capacity.

Experimental factors could also explain the lack of observed therapeutic effect. The rationale for applying RT prior to PRMT5 inhibition was to destroy all except TICs, enabling PRMT5 inhibition of TICs from which the tumor can recur. This strategy, which depends upon PRMT5 inhibition during the early stages of regrowth, may have failed because RT failed to completely destroy the proliferating tumor cells, or possibly because tumor regrowth had begun before PRMT5 inhibition was initiated. An alternative possibility is that the PRMT5 inhibitor did not sufficiently reduce PRMT5 activity to halt the development of tumor cells from the initiating stem-like cell population.

Further development of therapies targeting PRMT5 in PHGG might benefit from the development of more effective PRMT5 inhibitors or by combining PRMT5 inhibition with other means of regulating its activity. One possibility is to reduce the availability of the PRMT5 substrate S-adenosyl methionine (SAM), which is required for PRMT5 activity. This approach has been attempted successfully with other epigenetic regulatory drugs in preliminary preclinical experiments.^49^ A similar combinatory approach with a PRMT5 inhibitor could also be attempted.

The strengths of this study include the consistency of results across multiple cell lines, a mouse trial involving *PRMT5* KD, and ChIP-Seq in histone-mutant and non-mutant PHGG cell lines. The primary limitation is the inability to pharmacologically inhibit PRMT5 activity to a level that slows tumor growth.

## Acknowledgments

The Anschutz Medical Campus Functional Genomics Facility provided lentiviral-packaged shRNA constructs and the epigenetic shRNA library; the Anschutz Medical Campus Genomics Core performed bulk RNA sequencing of *PRMT5* KD and control samples and DNA sequencing of the ChIP-Seq samples for this project. BT245 cells were a gift from Keith Ligon; SU-DIPG-IV and SU-DIPG-XIII* cells were a gift from Michelle Monje; HSJD-DIPG-7 and HSJD-GBM-001 cells were a gift from Angel Carcaboso; VUMC-DIPG-10 cells were a gift from Esther Hulleman. Histological and immunocytochemistry staining was provided by the University of Colorado Cancer Center Pathology Shared Resource. The University of Colorado Anschutz Medical Campus Animal Imaging Shared Resource’s Small Animal Imaging Service provided MRI imaging of mice.

## Figure Captions

**Supplementary Figure 1.**
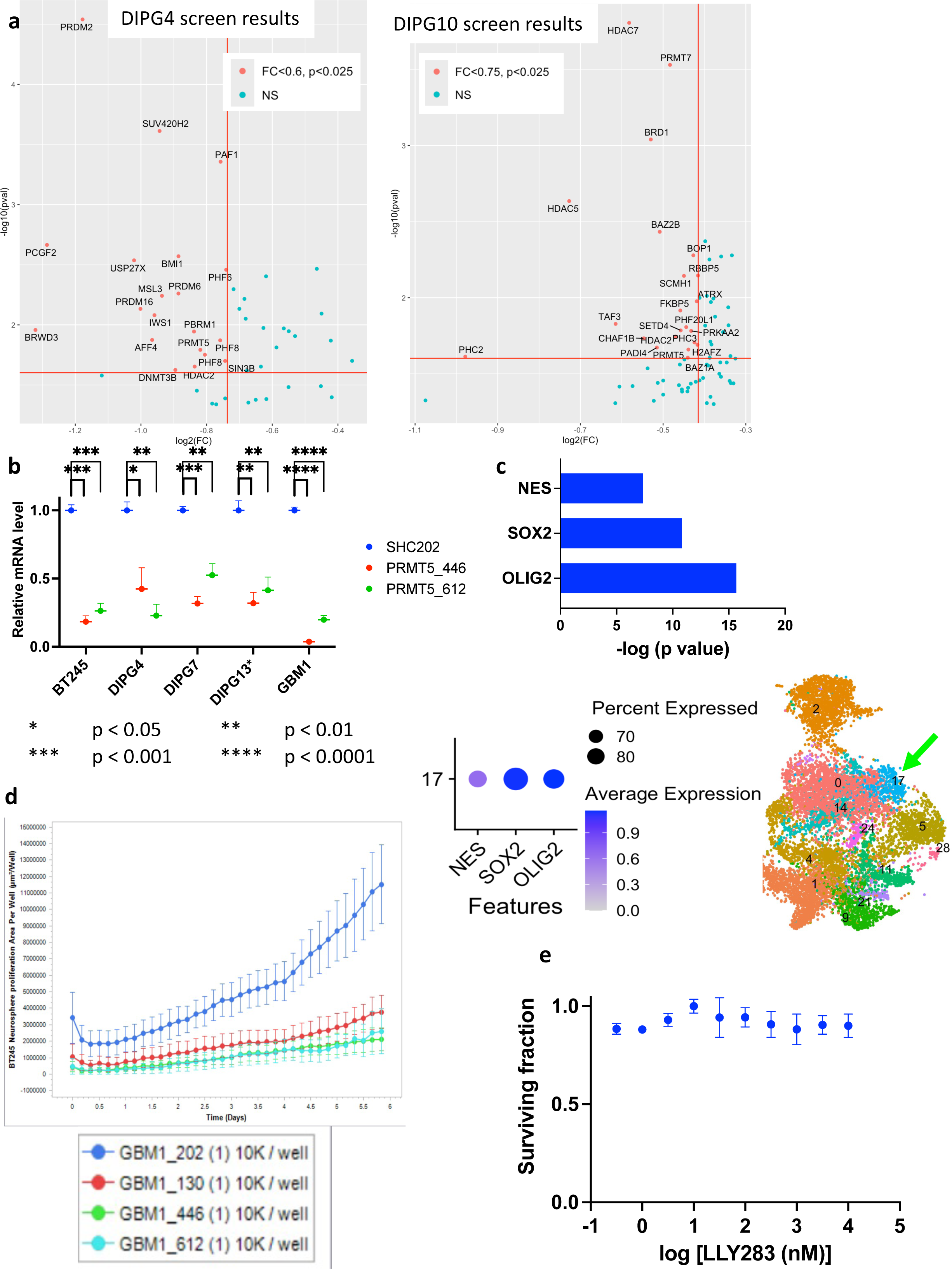
a. shRNA screen results showing genes important for cell growth in DIPG4 (left panel) and DIPG10 (right panel); significance thresholds are FC<0.6, p<0.025 for DIPG4 cells and FC<0.75, P<0.025 for DIPG10 cells; b. *PRMT5* KD effect on *PRMT5* expression in cell models of PHGG; c. single cell RNA-Seq analysis shows PHGG includes a population of stemlike cells that express stemness (*NES, SOX2*) and oligodendrocyte lineage (*OLIG2*) genes; upper panel shows –log p value of expression in stemlike population versus other tumor cell populations from a set of 19 PHGG patient samples; dot plot in lower left panel shows average differential expression and percentage of cells expressing each gene in stemlike population 17; lower right panel is a UMAP projection of cell types, arrowhead points to the stemlike population (17) depicted in upper panels; d. short-term cell growth assay of GBM1 cells cultured as neurospheres with *PRMT5* KD or empty vector control (blue curve – empty vector control; red, green and cyan curves – PRMT5 targeting shRNAs as shown in legend; error bars show SD); e. survival curve showing effect of LLY283 in normal human astrocyte cell line to demonstrate that PRMT5 inhibition using LLY283 has no effect on non-cancerous cells.

**Supplementary Figure 2.**
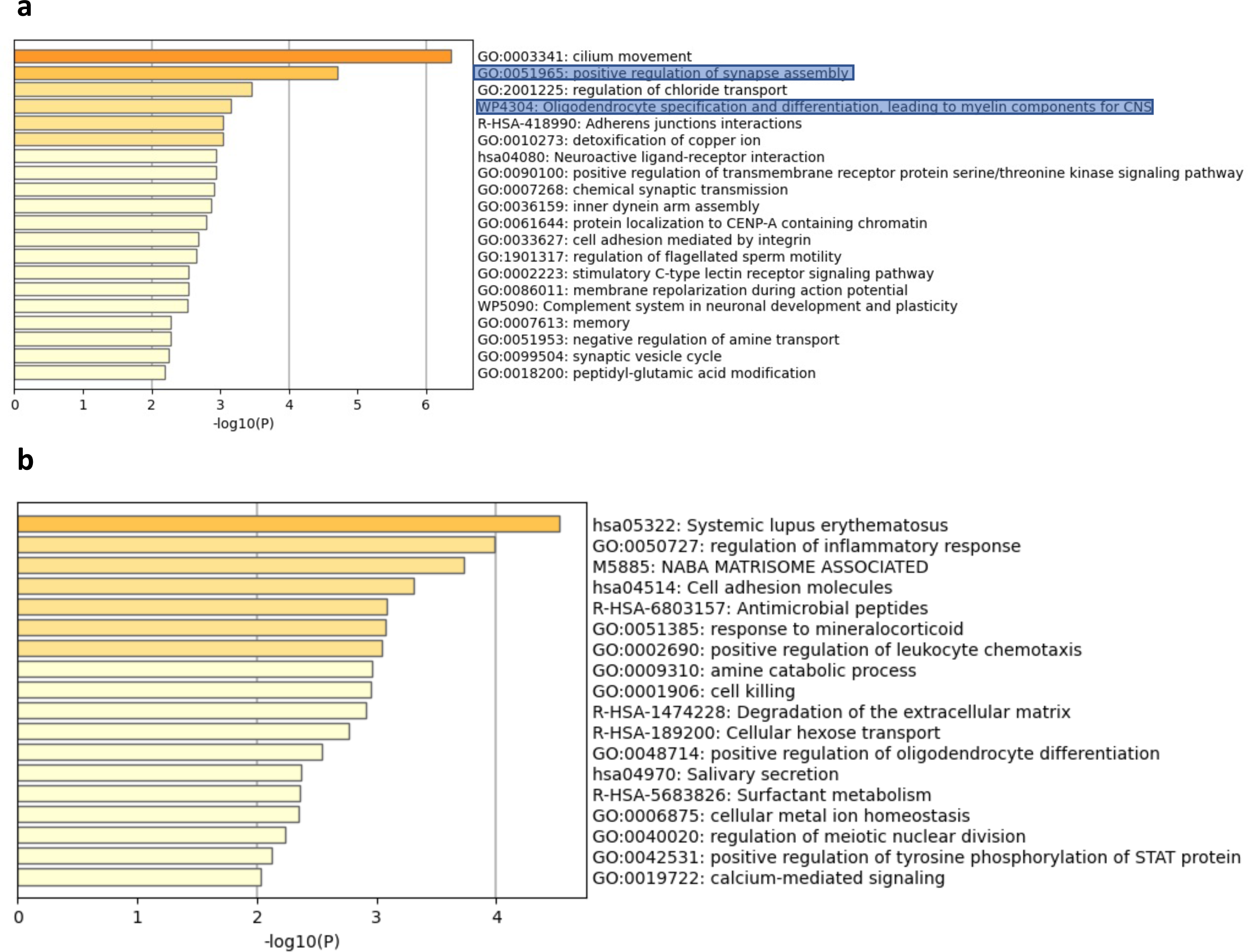
Metascape pathway analysis showed enrichment of differentiation pathways with *PRMT5* KD. a. Metascape analysis using 2,000 most differentially regulated genes with positive changes in log2 fold change as the result of *PRMT5* KD; b. Metascape analysis using 2,000 most differentially regulated genes with negative changes in log2 fold change as the result of *PRMT5* KD.

**Supplementary Figure 3.**
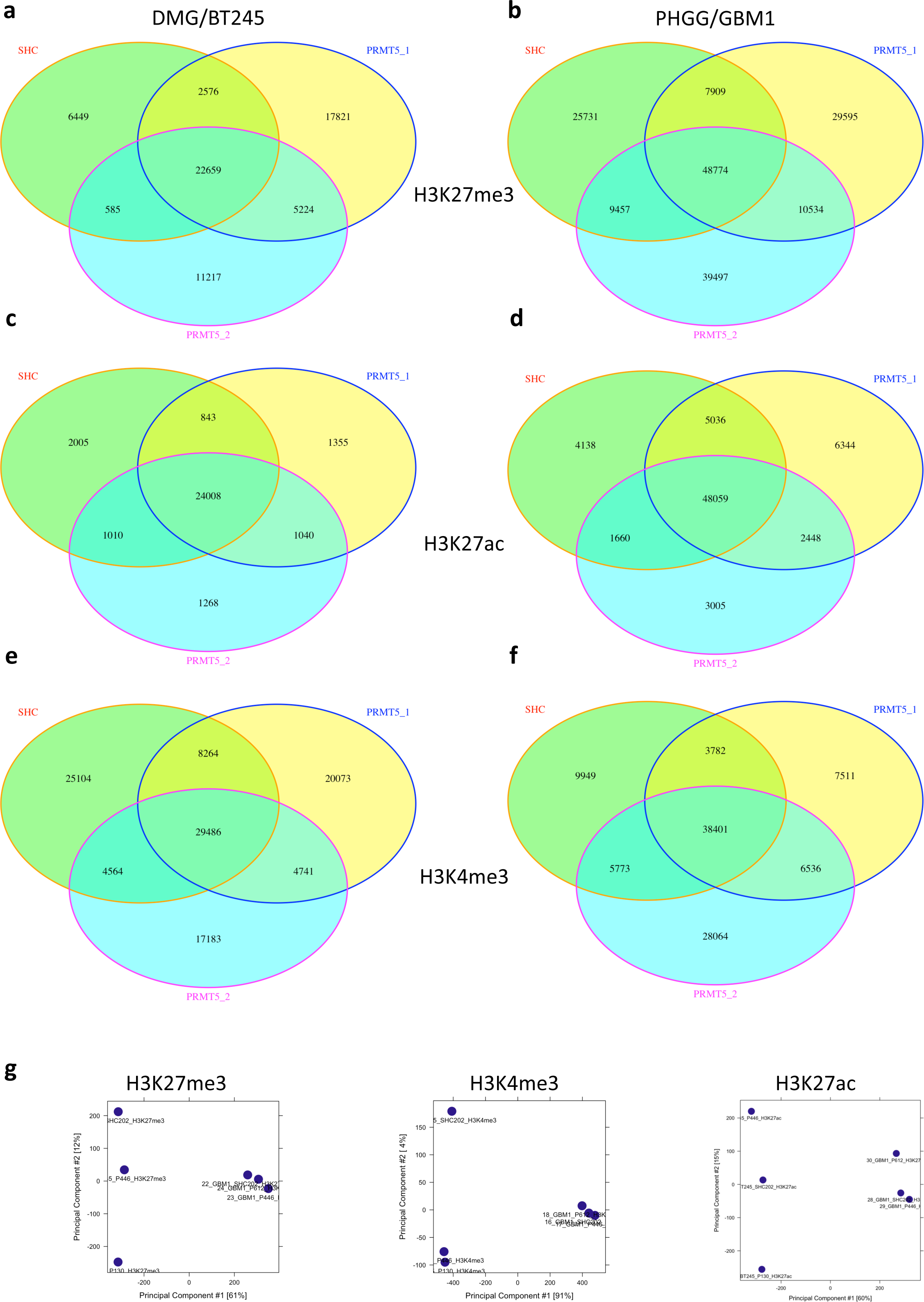
Overall effects of *PRMT5* KD on H3K27me3, H3K4me3 and H3K27ac occupancy in BT245 and GBM1 cells. a – b. Venn diagram showing overlap by genomic location of H3K27me3 occupancy in *PRMT5* KD (two samples) and empty vector control (one sample) in BT245 (panel a) and GBM1 (panel b) cells; regions of no overlap between *PRMT5* KD and empty vector control are input to differential expression analysis; c. – d. Venn diagram showing overlap by genomic location of H3K4me3 occupancy in *PRMT5* KD (two samples) and empty vector control (one sample) in BT245 (panel c) and GBM1 (panel d) cells; e. – f. Venn diagram showing overlap by genomic location of H3K27ac occupancy in *PRMT5* KD (two samples) and empty vector control (one sample) in BT245 (panel e) and GBM1 (panel f) cells; g. principal component plot of *PRMT5* KD (two samples per cell line) and empty vector control (one sample per cell line) based on differential occupancy of H3K27me3, H3K4me3 and H3K27ac at all genomic locations.

**Supplementary Figure 4.**
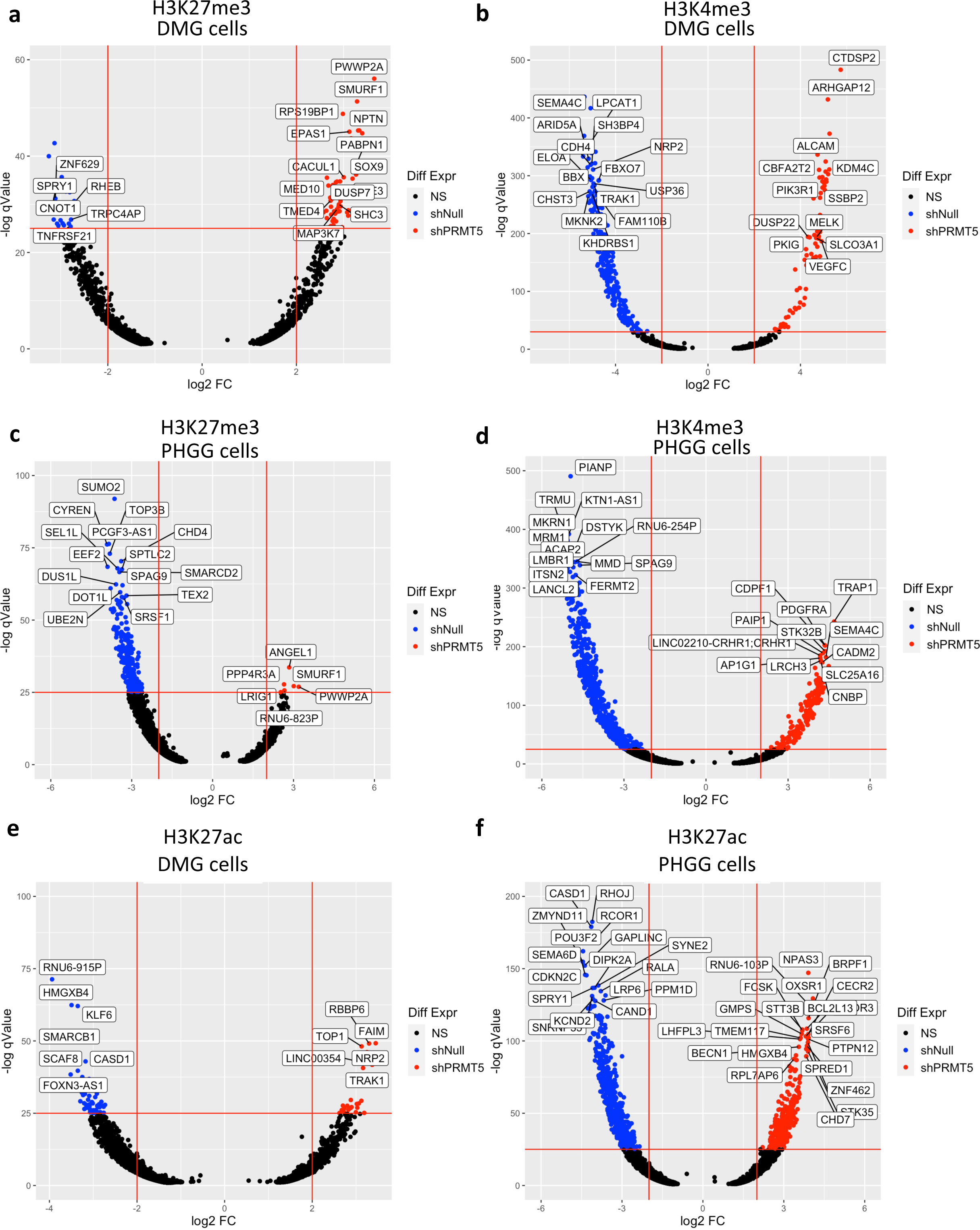
Differential effects of *PRMT5* KD on H3K27me3, H3K4me3 and H3K27ac occupancy in BT245 and GBM1 cells. a. BT245 cells, differential expression of H3K27me3 occupancy. Each point represents difference between occupancy in *PRMT5* KD versus control cells; genes of interest with |log2 FC| >2.0 and -log qValue >30 are labeled; b. BT245 cells, differential expression of H3K4me3 occupancy; c. GBM1 cells, differential expression of H3K27me3 occupancy; d. GBM1 cells, differential expression of H3K4me3 occupancy; differential expression analysis of H3K27ac occupancy at genes for BT245 (panel g) and GBM1 (panel h) cells (analysis is as described in Figure 6a-d).

**Supplementary Figure 5.**
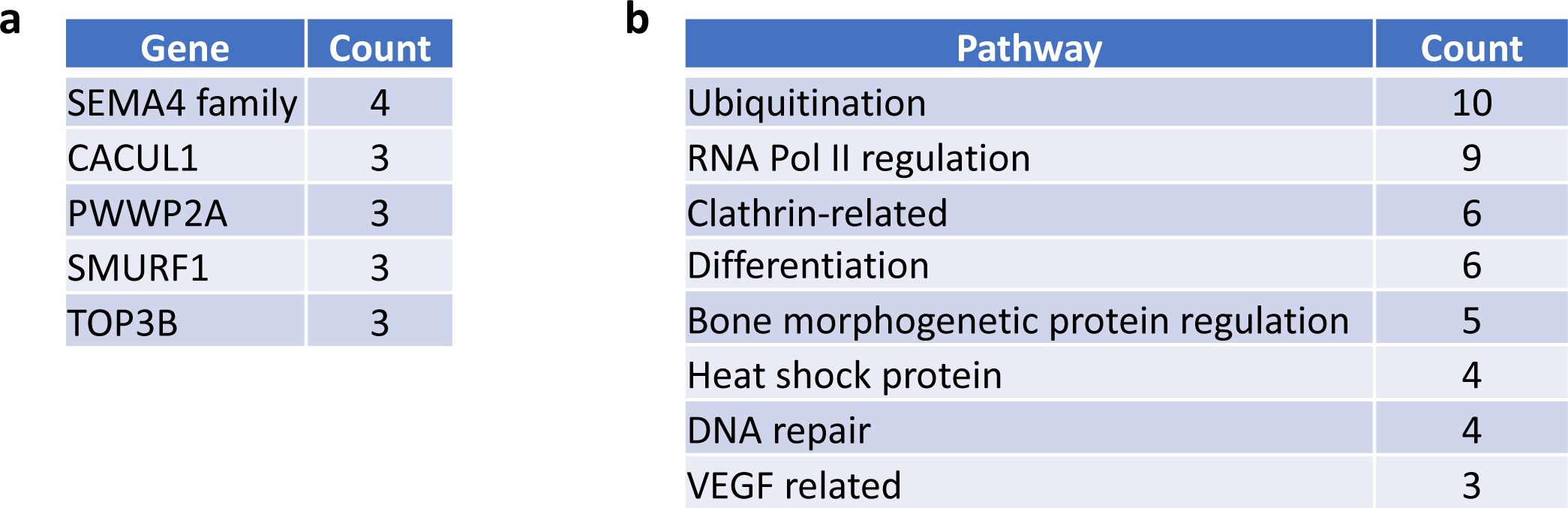
Summary of genes and pathways at which H3K4me3 or H3K27me3 were differentially expressed in *PRMT5* KD versus control cells. a. genes with differential expression; b. pathways controlled by genes listed in panel a.

**Supplementary Figure 6.**
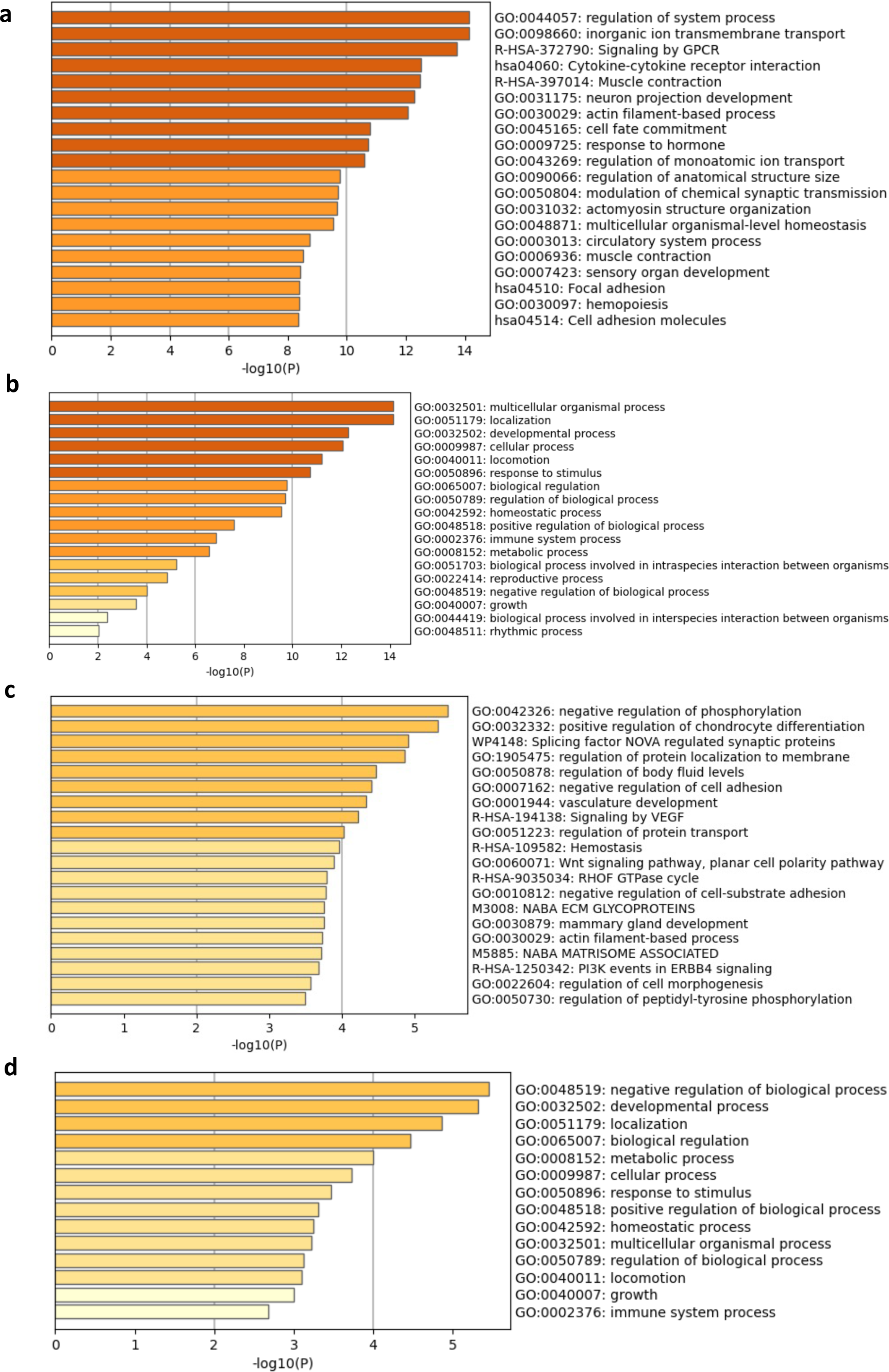
Metascape analysis of differential gene expression effects in BT245 cells. a. analysis of enrichment in gene ontology (GO) gene sets for H3K27me3 occupancy; b. analysis of enrichment in gene ontology – parent (GO-parent) gene sets for H3K27me3 occupancy; c. analysis of enrichment in GO gene sets for H3K4me3 occupancy; d. analysis of enrichment in GO-parent gene sets for H3K4me3 occupancy.

**Supplementary Figure 7.**
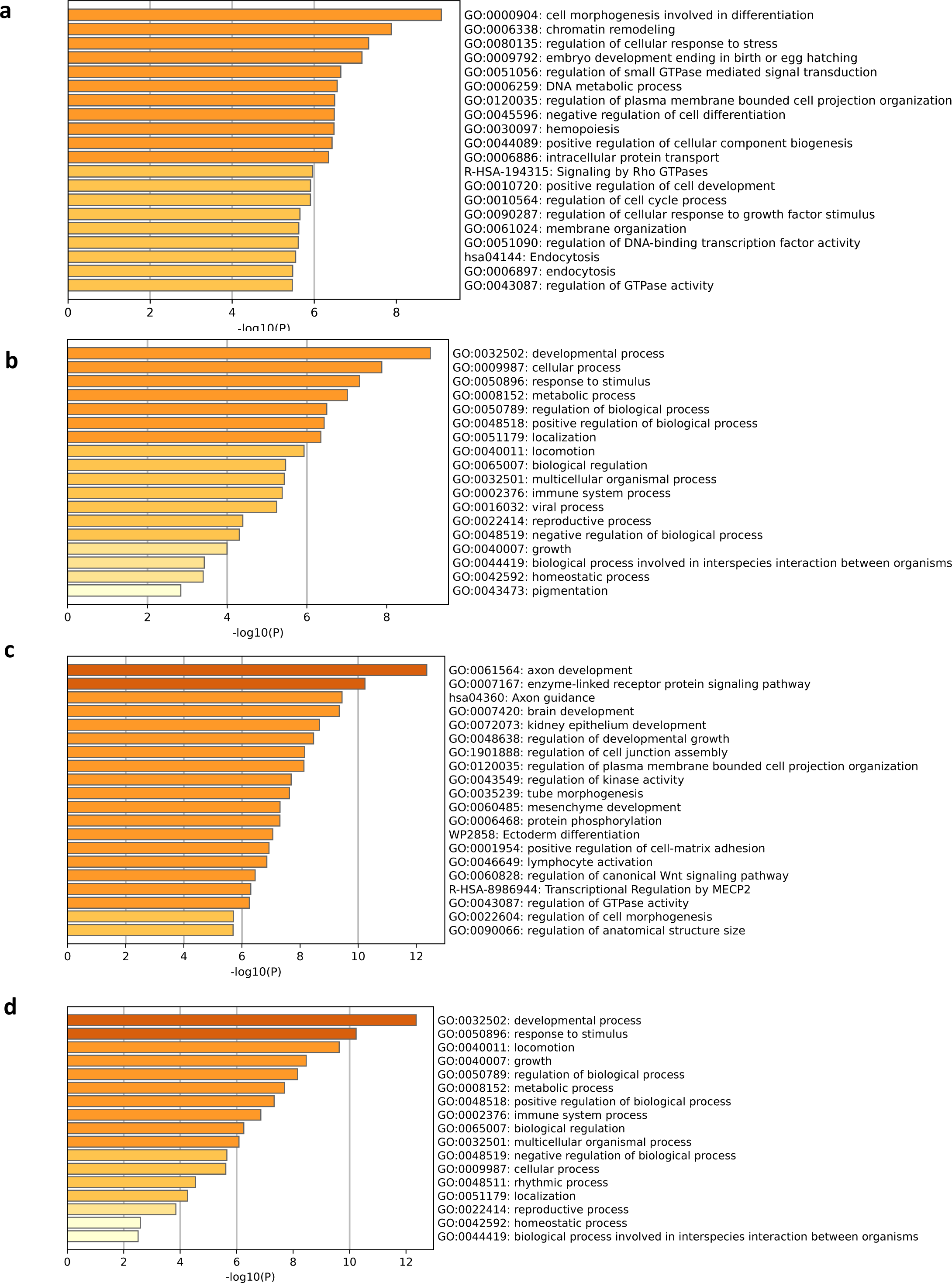
Metascape analysis of differential gene expression effects in GBM1 cells. a. analysis of enrichment in gene ontology (GO) gene sets for H3K27me3 occupancy; b. analysis of enrichment in gene ontology – parent (GO-parent) gene sets for H3K27me3 occupancy; c. analysis of enrichment in GO gene sets for H3K4me3 occupancy; d. analysis of enrichment in GO-parent gene sets for H3K4me3 occupancy.

