## Supplementary Tables 1 - 7 for "PRMT5 maintains tumor stem cells to promote pediatric high-grade glioma tumorigenesis"

**Supplementary Table 1   Cell line characteristics**

| Name | Type | Origin | Genetics | PDX |
|---|---|---|---|---|
| BT-245 | Thalamic DMG | Biopsy | H3.3K27M<br>*TP53* mutant | Y |
| SU-DIPG-IV (DIPG4) | Pontine DMG | Autopsy | H3.1K27M<br>*ACVR1* G328V | N |
| HSJD-DIPG-7 (DIPG7) | Pontine DMG | Autopsy | H3.3K27M<br>*ACVR1* R206H | N |
| VUMC-DIPG10 (DIPG10) | Pontine PHGG | Autopsy | H3K27 wt<br>NF1 Q209* | N |
| SU-DIPG-XIII (DIPG13) | Pontine DMG | Autopsy | H3.3K27M | Y |
| HSJD-GBM-001 (GBM1) | Cortical PHGG | Resection | | Y |

Supplementary Table 2 - Genes identified in shRNA screen of DIPG10 with FC<0.75 and p<0.025

| gene | FC | pval |
|---|---|---|
| HDAC7 | 0.66800347 | 0.00015638 |
| PRMT7 | 0.71533483 | 0.00029533 |
| BRD1 | 0.6926743 | 0.00091285 |
| HDAC5 | 0.60406447 | 0.00231881 |
| BAZ2B | 0.70309382 | 0.00368996 |
| HDAC6 | 0.75887868 | 0.00425791 |
| BOP1 | 0.74387452 | 0.00526863 |
| AURKB | 0.79352473 | 0.0052708 |
| PADI1 | 0.77937472 | 0.0053634 |
| SMARCA4 | 0.76417012 | 0.00560892 |
| RBBP5 | 0.74972215 | 0.00715111 |
| SCMH1 | 0.73226331 | 0.00718692 |
| ING2 | 0.7609046 | 0.00973224 |
| PRKCD | 0.751701 | 0.01001894 |
| EPC1 | 0.76924514 | 0.01013314 |
| ATRX | 0.74808909 | 0.01055675 |
| SATB1 | 0.76604744 | 0.01068763 |
| SIN3B | 0.7887178 | 0.01157343 |
| FKBP5 | 0.72777884 | 0.01213913 |
| SSRP1 | 0.7715067 | 0.01332264 |
| TAF3 | 0.65295961 | 0.01486317 |
| TDG | 0.78903922 | 0.01518298 |
| SUV39H1 | 0.75879651 | 0.01525792 |
| PHF20L1 | 0.73504749 | 0.01560617 |
| SETD4 | 0.72828902 | 0.01640594 |
| PRKAA2 | 0.74114833 | 0.01652201 |
| C14orf169 | 0.78193258 | 0.01683247 |
| HDAC2 | 0.72205449 | 0.01800291 |
| CHAF1B | 0.68462667 | 0.01850028 |
| CREBBP | 0.76551102 | 0.01945212 |
| PHC3 | 0.74525932 | 0.01967513 |
| KAT5 | 0.76681373 | 0.02030785 |
| H2AFZ | 0.7490171 | 0.02038748 |
| PADI4 | 0.69976412 | 0.02126194 |
| PRMT5 | 0.73768553 | 0.02191651 |
| SETD1B | 0.79160859 | 0.02418958 |
| PHC2 | 0.50749369 | 0.02438149 |
| BAZ1A | 0.73701683 | 0.02479513 |
| DNMT3A | 0.76452761 | 0.02483331 |
| KDM2A | 0.79818281 | 0.02511818 |
| UBE2I | 0.73289554 | 0.02547572 |

| | | |
|---|---|---|
| KDM1A | 0.77825105 | 0.02661978 |
| CHD1 | 0.78671284 | 0.02665045 |
| PHF21A | 0.72726818 | 0.02750327 |
| ATAD2B | 0.76087506 | 0.02752286 |
| ING4 | 0.76244649 | 0.02789743 |
| HDAC8 | 0.68895628 | 0.02912726 |
| PRDM10 | 0.7770081 | 0.03163104 |
| BRDT | 0.77918207 | 0.03308708 |
| ASXL2 | 0.75613798 | 0.03369588 |
| RPA3 | 0.75166605 | 0.03495237 |
| EZH2 | 0.70719131 | 0.03504187 |
| PRMT8 | 0.79054294 | 0.03523957 |
| ASH1L | 0.74686424 | 0.03569646 |
| CHD9 | 0.70076207 | 0.0362838 |
| NSD1 | 0.79207314 | 0.03639089 |
| RNF20 | 0.79567285 | 0.03652808 |
| TAF1L | 0.74072181 | 0.03672719 |
| CARM1 | 0.72612351 | 0.03757956 |
| USP22 | 0.67180063 | 0.0380942 |
| L3MBTL2 | 0.65719131 | 0.03838837 |
| ELP4 | 0.737382 | 0.03907479 |
| PHF8 | 0.72970337 | 0.04006104 |
| PBRM1 | 0.7365065 | 0.04129313 |
| SCML2 | 0.72995921 | 0.04139439 |
| POLR2B | 0.70718218 | 0.04335346 |
| KDM5D | 0.7655909 | 0.04480874 |
| KDM1B | 0.47471348 | 0.04730172 |
| TRIM28 | 0.69613512 | 0.0475829 |
| MECP2 | 0.76086591 | 0.04928561 |
| ING5 | 0.65265155 | 0.04933478 |
| SETMAR | 0.76970926 | 0.05000155 |

**Supplementary Table 3** Genes with p<0.05 when comparing expression FC between PRMT5 KD vs shNULL

| | BT245 | | DIPG4 | | DIPG7 | | DIPG13 | | GBM1 | |
|---|---|---|---|---|---|---|---|---|---|---|
| **gene** | **FC** | **p** | **FC** | **p** | **FC** | **p** | **FC** | **p** | **FC** | **p** |
| PAX3 | -0.39 | 0.0127 | -4.35 | 0.0014 | -4.66 | 0.0000 | 0.05 | 0.0348 | -2.22 | 0.0072 |
| PRMT5P1 | -2.66 | 0.0002 | -1.46 | 0.0001 | -1.48 | 0.0010 | -1.25 | 0.0000 | -1.59 | 0.0040 |
| MYO16 | -0.11 | 0.0001 | -1.36 | 0.0001 | -1.40 | 0.0001 | 0.60 | 0.0051 | -5.53 | 0.0005 |
| CDH4 | -0.59 | 0.0059 | -3.21 | 0.0003 | -4.57 | 0.0000 | 0.85 | 0.0013 | -0.12 | 0.0371 |
| CPNE7 | -0.34 | 0.0204 | -1.16 | 0.0016 | -3.17 | 0.0014 | 0.41 | 0.0492 | -3.13 | 0.0164 |
| PRMT5 | -1.64 | 0.0091 | -1.60 | 0.0016 | -1.14 | 0.0021 | -0.95 | 0.0395 | -1.90 | 0.0003 |
| KIF1A | -0.20 | 0.0284 | -3.16 | 0.0001 | -3.99 | 0.0000 | -0.05 | 0.0053 | 0.32 | 0.0181 |
| UCN2 | 0.89 | 0.0410 | -1.13 | 0.0105 | -3.52 | 0.0000 | -1.94 | 0.0426 | -1.25 | 0.0203 |
| MMP14 | -0.55 | 0.0029 | 0.18 | 0.0024 | -0.42 | 0.0182 | -1.64 | 0.0053 | -4.28 | 0.0000 |
| LINC01314 | -0.33 | 0.0237 | -2.08 | 0.0174 | -5.17 | 0.0000 | 1.13 | 0.0013 | 0.33 | 0.0120 |
| RHBDF2 | -0.69 | 0.0454 | 0.31 | 0.0015 | 0.28 | 0.0453 | -4.42 | 0.0000 | -1.02 | 0.0054 |
| ARPC1B | -0.46 | 0.0249 | 0.30 | 0.0004 | -0.28 | 0.0018 | -1.56 | 0.0059 | -3.37 | 0.0004 |
| CHL1 | -0.49 | 0.0494 | -2.72 | 0.0003 | -4.22 | 0.0004 | 0.86 | 0.0286 | 1.22 | 0.0298 |
| GAL3ST4 | -0.59 | 0.0154 | -1.67 | 0.0168 | -2.75 | 0.0000 | 0.46 | 0.0206 | 0.15 | 0.0117 |
| CNTNAP3P2 | -0.58 | 0.0107 | -1.73 | 0.0196 | -1.72 | 0.0452 | 0.48 | 0.0005 | -0.80 | 0.0302 |
| BACE2 | -0.02 | 0.0102 | -0.56 | 0.0260 | -2.17 | 0.0001 | -0.21 | 0.0064 | -1.34 | 0.0031 |
| MYADM | -0.45 | 0.0238 | -0.14 | 0.0051 | 0.19 | 0.0002 | -0.90 | 0.0160 | -2.56 | 0.0012 |
| ZFP3 | -1.31 | 0.0238 | 0.53 | 0.0006 | 0.89 | 0.0252 | 0.94 | 0.0058 | -3.91 | 0.0015 |
| DHRS4 | -0.18 | 0.0000 | 0.05 | 0.0255 | -3.02 | 0.0020 | 0.19 | 0.0012 | 0.21 | 0.0354 |
| CPM | 1.32 | 0.0392 | 0.75 | 0.0047 | -4.47 | 0.0005 | 1.70 | 0.0307 | -1.83 | 0.0157 |
| PLLP | 0.33 | 0.0158 | -1.44 | 0.0132 | -2.30 | 0.0022 | 0.85 | 0.0411 | 0.12 | 0.0290 |
| ENPP1 | -1.78 | 0.0472 | 0.61 | 0.0224 | 0.41 | 0.0122 | 1.08 | 0.0030 | -2.65 | 0.0188 |
| NLGN4X | -0.62 | 0.0031 | -2.70 | 0.0006 | -0.22 | 0.0237 | 1.04 | 0.0361 | 0.25 | 0.0023 |
| ARHGAP22 | -1.08 | 0.0165 | -0.52 | 0.0144 | -0.39 | 0.0377 | 0.24 | 0.0031 | -0.43 | 0.0358 |
| COL4A1 | -0.31 | 0.0487 | 0.21 | 0.0082 | 0.35 | 0.0302 | -1.58 | 0.0000 | -0.65 | 0.0008 |
| XPO7 | -0.35 | 0.0003 | -0.17 | 0.0019 | -0.29 | 0.0136 | -0.48 | 0.0122 | -0.20 | 0.0140 |
| BNIP3P5 | -0.44 | 0.0228 | 0.86 | 0.0013 | -0.87 | 0.0358 | -1.62 | 0.0352 | 0.67 | 0.0046 |
| PAQR8 | -0.19 | 0.0001 | -0.60 | 0.0138 | -0.96 | 0.0287 | 0.62 | 0.0003 | -0.19 | 0.0448 |
| DAP | -0.24 | 0.0464 | 0.10 | 0.0051 | -0.95 | 0.0003 | -0.39 | 0.0096 | 0.18 | 0.0127 |
| FTH1P5 | -0.53 | 0.0127 | 0.47 | 0.0002 | -0.52 | 0.0212 | -1.51 | 0.0008 | 0.79 | 0.0012 |
| KIRREL | -0.05 | 0.0012 | 0.31 | 0.0113 | 0.20 | 0.0412 | -0.69 | 0.0025 | -0.76 | 0.0150 |
| SOD2 | 0.02 | 0.0001 | 0.47 | 0.0029 | -0.29 | 0.0104 | -0.90 | 0.0083 | -0.14 | 0.0419 |
| KDM5C | -0.22 | 0.0175 | 0.18 | 0.0078 | 0.13 | 0.0159 | -0.72 | 0.0003 | -0.03 | 0.0488 |
| ELP5 | 0.46 | 0.0004 | -0.23 | 0.0082 | -0.36 | 0.0169 | -0.42 | 0.0003 | 0.02 | 0.0003 |
| ANXA6 | -0.27 | 0.0452 | 0.10 | 0.0000 | -0.18 | 0.0417 | 0.33 | 0.0106 | -0.12 | 0.0270 |
| MIOS | -0.51 | 0.0174 | 0.04 | 0.0114 | 0.31 | 0.0005 | 0.32 | 0.0162 | -0.20 | 0.0177 |
| MRPL36 | 0.36 | 0.0078 | -0.14 | 0.0234 | -0.58 | 0.0354 | 0.56 | 0.0003 | -0.16 | 0.0351 |
| RNF139 | 0.42 | 0.0019 | -0.05 | 0.0043 | 0.47 | 0.0354 | -0.28 | 0.0037 | -0.34 | 0.0061 |
| CTC-479C5.1( | -0.19 | 0.0001 | -0.13 | 0.0012 | 1.27 | 0.0044 | -1.01 | 0.0082 | 0.40 | 0.0397 |
| SYVN1 | 0.10 | 0.0472 | 0.39 | 0.0003 | 0.51 | 0.0097 | -1.01 | 0.0059 | 0.37 | 0.0470 |
| C1QTNF3 | -0.39 | 0.0172 | -0.38 | 0.0073 | -1.36 | 0.0006 | 1.76 | 0.0418 | 1.00 | 0.0429 |
| RSBN1L | 0.14 | 0.0001 | 0.28 | 0.0345 | 0.22 | 0.0298 | 0.31 | 0.0250 | 0.04 | 0.0293 |

| | | | | | | | | | | |
|---|---|---|---|---|---|---|---|---|---|---|
| VAMP3 | 0.66 | 0.0384 | 0.29 | 0.0166 | 0.45 | 0.0036 | -0.42 | 0.0007 | 0.23 | 0.0021 |
| PIK3R1 | 0.36 | 0.0136 | 0.17 | 0.0056 | -0.40 | 0.0038 | 0.82 | 0.0007 | 0.31 | 0.0110 |
| NOD1 | 0.50 | 0.0191 | 0.29 | 0.0090 | 0.68 | 0.0002 | 0.36 | 0.0336 | -0.53 | 0.0002 |
| ZNF182 | 0.52 | 0.0031 | 0.27 | 0.0063 | 0.30 | 0.0287 | 0.26 | 0.0035 | 0.17 | 0.0017 |
| CTD-2015B23 | 0.92 | 0.0417 | 0.42 | 0.0466 | -0.98 | 0.0355 | 0.63 | 0.0023 | 0.55 | 0.0350 |
| KIAA0408 | 1.15 | 0.0059 | -0.60 | 0.0115 | 0.31 | 0.0143 | 0.51 | 0.0138 | 0.49 | 0.0033 |
| PUS7 | 0.61 | 0.0337 | 0.14 | 0.0143 | 0.30 | 0.0059 | 0.95 | 0.0109 | -0.15 | 0.0061 |
| RP11-244H3.4 | 22.91 | 0.0000 | 1.22 | 0.0336 | 10.02 | 0.0320 | -0.15 | 0.0239 | -0.97 | 0.0199 |

**Supplementary Table 4** BT245 H3K4me3 GSEA gene sets containing target genes

| | NAME | SIZE | NES | QVAL |
|---|---|---|---|---|
| 1 | ZNF250_TARGET_GENES | 30 | -2.581579 | 0.00044794 |
| 2 | RAG1_TARGET_GENES | 73 | -2.5407476 | 0.00047803 |
| 3 | SETD1A_TARGET_GENES | 73 | -2.5301251 | 0.0003718 |
| 4 | GARY_CD5_TARGETS_UP | 38 | -2.4810898 | 0.00079551 |
| 5 | ZNF224_TARGET_GENES | 99 | -2.4340396 | 0.00156404 |
| 6 | NAB2_TARGET_GENES | 130 | -2.3409944 | 0.00297364 |
| 7 | ZBED5_TARGET_GENES | 129 | -2.3377926 | 0.00308108 |
| 8 | ZNF610_TARGET_GENES | 51 | -2.2863123 | 0.00409315 |
| 9 | ZZZ3_TARGET_GENES | 17 | -2.236395 | 0.00659592 |
| 10 | ZNF711_TARGET_GENES | 97 | -2.218405 | 0.0070183 |
| 11 | KLF7_TARGET_GENES | 56 | -2.2120678 | 0.0071241 |
| 12 | ZSCAN31_TARGET_GENES | 35 | -2.1914692 | 0.00775869 |
| 13 | ZNF184_TARGET_GENES | 98 | -2.1865716 | 0.00793026 |
| 14 | ZNF423_TARGET_GENES | 101 | -2.1831098 | 0.00779958 |
| 15 | ZNF260_TARGET_GENES | 45 | -2.170383 | 0.00800798 |
| 16 | SENESE_HDAC3_TARGETS_DN | 47 | -2.1668353 | 0.00802506 |
| 17 | PAX6_TARGET_GENES | 59 | -2.1636512 | 0.00791087 |
| 18 | ZNF592_TARGET_GENES | 146 | -2.157691 | 0.00807314 |
| 19 | TAKEDA_TARGETS_OF_NUP98_HOXA9_FUSION_3D_UP | 21 | -2.1449754 | 0.00880833 |
| 20 | ZNF92_TARGET_GENES | 138 | -2.1337247 | 0.00955016 |
| 21 | RBM34_TARGET_GENES | 92 | -2.1303046 | 0.00982597 |
| 22 | FISCHER_DREAM_TARGETS | 68 | -2.1289954 | 0.00993578 |
| 23 | ADA2_TARGET_GENES | 54 | -2.1220427 | 0.01026205 |
| 24 | DYRK1A_TARGET_GENES | 39 | -2.1146972 | 0.01073436 |
| 25 | DIDO1_TARGET_GENES | 80 | -2.111983 | 0.01091244 |
| 26 | KAT2A_TARGET_GENES | 53 | -2.1027555 | 0.01155392 |
| 27 | ARID5B_TARGET_GENES | 65 | -2.1003673 | 0.01153221 |
| 28 | POU2AF1_TARGET_GENES | 80 | -2.1001537 | 0.01152504 |
| 29 | HOXA1_TARGET_GENES | 60 | -2.097193 | 0.01172225 |
| 30 | ZNF843_TARGET_GENES | 87 | -2.0961869 | 0.0117591 |
| 31 | ZNF2_TARGET_GENES | 88 | -2.094914 | 0.01182673 |
| 32 | JOHNSTONE_PARVB_TARGETS_3_UP | 49 | -2.082333 | 0.0125687 |
| 33 | CDC5L_TARGET_GENES | 26 | -2.0815456 | 0.01255634 |
| 34 | SKIL_TARGET_GENES | 139 | -2.0795658 | 0.01249339 |
| 35 | ZNF528_TARGET_GENES | 54 | -2.0790193 | 0.01248301 |
| 36 | TET1_TARGET_GENES | 78 | -2.0657282 | 0.01376465 |
| 37 | HMG20B_TARGET_GENES | 146 | -2.0531657 | 0.01488688 |
| 38 | BARX1_TARGET_GENES | 84 | -2.046084 | 0.01552011 |
| 39 | ZNF30_TARGET_GENES | 91 | -2.0374367 | 0.0163697 |
| 40 | SS18_SSX1_FUSION_UNIPROT_Q8IZH1_UNREVIEWED_TARGET | 46 | -2.036557 | 0.0164597 |
| 41 | SMN1_SMN2_TARGET_GENES | 63 | -2.0282764 | 0.01736218 |

| | | | | |
|---|---|---|---|---|
| 42 | ZNF597_TARGET_GENES | 71 | -2.027818 | 0.01733137 |
| 43 | ZNF407_TARGET_GENES | 91 | -2.0258725 | 0.01756246 |
| 44 | PEDRIOLI_MIR31_TARGETS_DN | 47 | -2.0218024 | 0.01767615 |
| 45 | WANG_LMO4_TARGETS_UP | 39 | -1.9999914 | 0.02035257 |
| 46 | BASAKI_YBX1_TARGETS_DN | 38 | -1.9955444 | 0.02085539 |
| 47 | FEV_TARGET_GENES | 94 | -1.9926442 | 0.02103882 |
| 48 | EPC1_TARGET_GENES | 26 | -1.991858 | 0.02105329 |
| 49 | ZBTB5_TARGET_GENES | 35 | -1.9911547 | 0.02111963 |
| 50 | SFMBT1_TARGET_GENES | 126 | -1.989852 | 0.02127123 |
| 51 | ZFP91_TARGET_GENES | 125 | -1.9847974 | 0.02211872 |
| 52 | TERF2_TARGET_GENES | 16 | -1.9836761 | 0.02208694 |
| 53 | TBX3_TARGET_GENES | 50 | -1.9832629 | 0.0220909 |
| 54 | LOPEZ_MBD_TARGETS | 78 | -1.9743214 | 0.02293245 |
| 55 | CDC73_TARGET_GENES | 36 | -1.9702588 | 0.02355747 |
| 56 | YAMAZAKI_TCEB3_TARGETS_DN | 23 | -1.9653615 | 0.02405516 |
| 57 | FOXN3_TARGET_GENES | 124 | -1.9651221 | 0.02397391 |
| 58 | ID1_TARGET_GENES | 97 | -1.9589885 | 0.02478412 |
| 59 | ARNT2_TARGET_GENES | 42 | -1.9581004 | 0.02486458 |
| 60 | WIERENGA_STAT5A_TARGETS_UP | 26 | -1.9572781 | 0.02485676 |
| 61 | ZNF320_TARGET_GENES | 116 | -1.9496479 | 0.02610237 |
| 62 | ZSCAN30_TARGET_GENES | 137 | -1.9452657 | 0.02669576 |
| 63 | SNRNP70_TARGET_GENES | 46 | -1.94159 | 0.02704641 |
| 64 | HAND1_TARGET_GENES | 48 | -1.9303129 | 0.02869524 |
| 65 | SANSOM_APC_MYC_TARGETS | 22 | -1.9300749 | 0.02868338 |
| 66 | HSD17B8_TARGET_GENES | 41 | -1.9269478 | 0.02914829 |
| 67 | ZNF549_TARGET_GENES | 89 | -1.9266835 | 0.02907563 |
| 68 | SNIP1_TARGET_GENES | 45 | -1.926063 | 0.02918754 |
| 69 | LEI_MYB_TARGETS | 20 | -1.921667 | 0.02997301 |
| 70 | PRKDC_TARGET_GENES | 55 | -1.9213849 | 0.03001028 |
| 71 | IRF5_TARGET_GENES | 45 | -1.9184301 | 0.03038422 |
| 72 | FORTSCHEGGER_PHF8_TARGETS_DN | 81 | -1.9155843 | 0.0307246 |
| 73 | KIM_WT1_TARGETS_12HR_UP | 20 | -1.9116365 | 0.03147934 |
| 74 | CREB3_TARGET_GENES | 42 | -1.9109308 | 0.03153053 |
| 75 | MZF1_TARGET_GENES | 130 | -1.9022454 | 0.03306616 |
| 76 | HHEX_TARGET_GENES | 63 | -1.8985568 | 0.03370873 |
| 77 | ZNF589_TARGET_GENES | 51 | -1.8961159 | 0.03403033 |
| 78 | ZNF322_TARGET_GENES | 67 | -1.8924718 | 0.03454669 |
| 79 | ZNF768_TARGET_GENES | 111 | -1.8916018 | 0.03474388 |
| 80 | ZNF282_TARGET_GENES | 83 | -1.8864549 | 0.03535264 |
| 81 | NUYTTEN_NIPP1_TARGETS_UP | 69 | -1.8796265 | 0.03652843 |
| 82 | GAUSSMANN_MLL_AF4_FUSION_TARGETS_F_UP | 25 | -1.8789607 | 0.036625 |
| 83 | NFKBIA_TARGET_GENES | 105 | -1.8781006 | 0.03669048 |
| 84 | ZNF318_TARGET_GENES | 89 | -1.8712276 | 0.03807587 |

| | | | | |
|---|---|---|---|---|
| 85 | SHEN_SMARCA2_TARGETS_UP | 26 | -1.8709605 | 0.03810007 |
| 86 | ADNP_TARGET_GENES | 52 | -1.8692873 | 0.03818058 |
| 87 | ID2_TARGET_GENES | 64 | -1.8652823 | 0.03870332 |
| 88 | UDAYAKUMAR_MED1_TARGETS_DN | 15 | -1.8636485 | 0.03890031 |
| 89 | CBX7_TARGET_GENES | 81 | -1.8627263 | 0.03888126 |
| 90 | MYOCD_TARGET_GENES | 125 | -1.8623333 | 0.03882173 |
| 91 | BARX2_TARGET_GENES | 97 | -1.861858 | 0.03890801 |
| 92 | FOXD2_TARGET_GENES | 79 | -1.8582191 | 0.03964126 |
| 93 | CC2D1A_TARGET_GENES | 117 | -1.853242 | 0.0407477 |
| 94 | ZNF391_TARGET_GENES | 61 | -1.8380694 | 0.04388569 |
| 95 | ZNF436_TARGET_GENES | 50 | -1.8352838 | 0.04433072 |
| 96 | NPM1_TARGET_GENES | 21 | -1.8287069 | 0.04564423 |
| 97 | GARY_CD5_TARGETS_DN | 27 | -1.8267695 | 0.04578838 |
| 98 | NUYTTEN_EZH2_TARGETS_DN | 61 | -1.826642 | 0.0457648 |
| 99 | TAFAZZIN_TARGET_GENES | 48 | -1.8261557 | 0.04574166 |
| 100 | KAT5_TARGET_GENES | 37 | -1.8229932 | 0.04615664 |
| 101 | NR1H4_TARGET_GENES | 30 | -1.8186346 | 0.04736222 |
| 102 | ZNF10_TARGET_GENES | 55 | -1.8186144 | 0.04730268 |
| 103 | ZNF547_TARGET_GENES | 17 | -1.8156589 | 0.0478083 |
| 104 | ZBTB12_TARGET_GENES | 74 | -1.8152723 | 0.04775319 |
| 105 | BARHL1_TARGET_GENES | 63 | -1.8134491 | 0.04823965 |
| 106 | ZNF394_TARGET_GENES | 69 | -1.8105391 | 0.04885524 |

**Supplementary Table 5** BT245 H3K27me3 GSEA gene sets containing target genes

| | NAME | SIZE | NES | QVAL |
|---|---|---|---|---|
| 1 | ZFP91_TARGET_GENES | 234 | -3.001304 | 0.00097389 |
| 2 | SUPT20H_TARGET_GENES | 131 | -2.9542046 | 0.00143943 |
| 3 | ZSCAN30_TARGET_GENES | 267 | -2.8551836 | 0.00419532 |
| 4 | ZNF184_TARGET_GENES | 220 | -2.7898235 | 0.00322215 |
| 5 | NKX2_2_TARGET_GENES | 149 | -2.757969 | 0.00359064 |
| 6 | RFX7_TARGET_GENES | 58 | -2.7432728 | 0.00332989 |
| 7 | ZBED5_TARGET_GENES | 219 | -2.7412505 | 0.0029969 |
| 8 | ZNF34_TARGET_GENES | 63 | -2.7265403 | 0.00252699 |
| 9 | ZNF660_TARGET_GENES | 105 | -2.712267 | 0.0027608 |
| 10 | KINSEY_TARGETS_OF_EWSR1_FLII_FUSION_UP | 190 | -2.6436286 | 0.00462046 |
| 11 | LOPEZ_MBD_TARGETS | 168 | -2.6271808 | 0.00488608 |
| 12 | ZNF547_TARGET_GENES | 52 | -2.6134996 | 0.00523643 |
| 13 | AHRR_TARGET_GENES | 153 | -2.5912242 | 0.00643646 |
| 14 | BARHL1_TARGET_GENES | 130 | -2.56462 | 0.00698693 |
| 15 | CEBPZ_TARGET_GENES | 150 | -2.5207958 | 0.00704199 |
| 16 | ZC3H11A_TARGET_GENES | 118 | -2.4846208 | 0.00892084 |
| 17 | DIDO1_TARGET_GENES | 165 | -2.4655108 | 0.00943172 |
| 18 | BENPORATH_MYC_MAX_TARGETS | 81 | -2.4309895 | 0.01177904 |
| 19 | ZNF592_TARGET_GENES | 297 | -2.4309473 | 0.0115687 |
| 20 | HDGF_TARGET_GENES | 149 | -2.3947716 | 0.01284928 |
| 21 | ELF2_TARGET_GENES | 126 | -2.3813453 | 0.01376294 |
| 22 | SETD1A_TARGET_GENES | 130 | -2.3580718 | 0.01510077 |
| 23 | DBP_TARGET_GENES | 145 | -2.3318355 | 0.01717624 |
| 24 | ZNF740_TARGET_GENES | 223 | -2.3099914 | 0.01956846 |
| 25 | SUMO1_TARGET_GENES | 126 | -2.2726092 | 0.0237773 |
| 26 | ADA2_TARGET_GENES | 108 | -2.246792 | 0.02670675 |
| 27 | KAT2A_TARGET_GENES | 120 | -2.2442462 | 0.02696402 |
| 28 | BENPORATH_SOX2_TARGETS | 112 | -2.2389789 | 0.02725508 |
| 29 | ZNF711_TARGET_GENES | 167 | -2.2302682 | 0.02782062 |
| 30 | NFE2L1_TARGET_GENES | 235 | -2.2156193 | 0.02979411 |
| 31 | ZBTB18_TARGET_GENES | 107 | -2.2152786 | 0.02941958 |
| 32 | ATF6_TARGET_GENES | 104 | -2.2108076 | 0.02990539 |
| 33 | NUYTTEN_EZH2_TARGETS_DN | 129 | -2.2059968 | 0.03028427 |
| 34 | TEAD2_TARGET_GENES | 172 | -2.1959548 | 0.03191398 |
| 35 | E2F2_TARGET_GENES | 170 | -2.1931462 | 0.03230707 |
| 36 | GLI3_TARGET_GENES | 79 | -2.189618 | 0.03227998 |
| 37 | ZNF30_TARGET_GENES | 191 | -2.1833298 | 0.03365453 |
| 38 | SNIP1_TARGET_GENES | 82 | -2.180672 | 0.03363878 |
| 39 | DLX6_TARGET_GENES | 66 | -2.174071 | 0.03427268 |
| 40 | ZNF213_TARGET_GENES | 76 | -2.1653056 | 0.03485603 |
| 41 | BENPORATH_NANOG_TARGETS | 151 | -2.1650531 | 0.0346734 |

| | | | | |
|---|---|---|---|---|
| 42 | ZNF664_TARGET_GENES | 189 | -2.1616006 | 0.03519701 |
| 43 | ZNF618_TARGET_GENES | 217 | -2.1615229 | 0.03478292 |
| 44 | NR1I2_TARGET_GENES | 38 | -2.1564248 | 0.03501825 |
| 45 | ZNF8_TARGET_GENES | 124 | -2.1532826 | 0.03547489 |
| 46 | CREB3L4_TARGET_GENES | 151 | -2.1419082 | 0.03651228 |
| 47 | TET1_TARGET_GENES | 146 | -2.1414459 | 0.03606115 |
| 48 | CASP8AP2_TARGET_GENES | 40 | -2.1344557 | 0.03706257 |
| 49 | ZNF85_TARGET_GENES | 33 | -2.1329443 | 0.03723522 |
| 50 | ZNF391_TARGET_GENES | 126 | -2.1181602 | 0.0380046 |
| 51 | ASH1L_TARGET_GENES | 145 | -2.1158276 | 0.0380563 |
| 52 | ZFHX3_TARGET_GENES | 193 | -2.1111116 | 0.03723473 |
| 53 | ZSCAN5DP_TARGET_GENES | 173 | -2.1041086 | 0.03871692 |
| 54 | ZNF2_TARGET_GENES | 184 | -2.1016161 | 0.03885844 |
| 55 | ZNF318_TARGET_GENES | 188 | -2.0752702 | 0.04177542 |
| 56 | JOHNSTONE_PARVB_TARGETS_2_DN | 59 | -2.0667543 | 0.04357056 |
| 57 | ZNF257_TARGET_GENES | 83 | -2.05676 | 0.04408435 |
| 58 | SHEN_SMARCA2_TARGETS_UP | 62 | -2.0549047 | 0.04446285 |
| 59 | WRNIP1_TARGET_GENES | 134 | -2.0527117 | 0.04458369 |
| 60 | ZNF697_TARGET_GENES | 43 | -2.0381796 | 0.04686742 |
| 61 | RBM34_TARGET_GENES | 134 | -2.026903 | 0.0484175 |
| 62 | JOHNSTONE_PARVB_TARGETS_3_DN | 106 | 2.8805237 | 0.00120939 |
| 63 | BARX1_TARGET_GENES | 133 | 2.5018966 | 0.02330758 |
| 64 | ZBTB12_TARGET_GENES | 154 | 2.4702668 | 0.02730962 |
| 65 | ZNF507_TARGET_GENES | 85 | 2.392494 | 0.03777168 |
| 66 | BARX2_TARGET_GENES | 174 | 2.376268 | 0.03809072 |
| 67 | CC2D1A_TARGET_GENES | 234 | 2.2959542 | 0.0451983 |
| 68 | SALL4_TARGET_GENES | 206 | 2.2900512 | 0.04293304 |
| 69 | CIITA_TARGET_GENES | 122 | 2.2818937 | 0.04353983 |
| 70 | JACKSON_DNMT1_TARGETS_UP | 19 | 2.2533557 | 0.04646672 |

**Supplementary Table 6** GBM1 H3K4me3 GSEA gene sets containing target genes

| | NAME | SIZE | NES | QVAL |
|---|---|---|---|---|
| 1 | KIM_MYCN_AMPLIFICATION_TARGETS_DN | 17 | -2.4501052 | 0.00539568 |
| 2 | VANLOO_SP3_TARGETS_DN | 19 | -2.283598 | 0.00785038 |
| 3 | KOINUMA_TARGETS_OF_SMAD2_OR_SMAD3 | 136 | -2.2609813 | 0.00811342 |
| 4 | YAMAZAKI_TCEB3_TARGETS_UP | 33 | -2.2474935 | 0.00771605 |
| 5 | LU_EZH2_TARGETS_DN | 59 | -2.240434 | 0.00779646 |
| 6 | CUI_TCF21_TARGETS_2_DN | 185 | -2.226008 | 0.00834097 |
| 7 | RIEGE_DELTANP63_DIRECT_TARGETS_UP | 26 | -2.2186081 | 0.00890989 |
| 8 | GUO_HEX_TARGETS_UP | 18 | -2.189631 | 0.00952796 |
| 9 | KINSEY_TARGETS_OF_EWSR1_FLII_FUSION_UP | 202 | -2.0600042 | 0.02030207 |
| 10 | MCBRYAN_PUBERTAL_TGFB1_TARGETS_UP | 36 | -2.03804 | 0.02288465 |
| 11 | LU_EZH2_TARGETS_UP | 33 | -1.9782336 | 0.02884014 |
| 12 | AFFAR_YY1_TARGETS_UP | 35 | -1.9777827 | 0.02885721 |
| 13 | KATSANOU_ELAVL1_TARGETS_UP | 42 | -1.976529 | 0.02882184 |
| 14 | PEDRIOLI_MIR31_TARGETS_UP | 40 | -1.9736574 | 0.0290944 |
| 15 | TONKS_TARGETS_OF_RUNX1_RUNX1T1_FUSION_HSC_UP | 34 | -1.9728575 | 0.02894134 |
| 16 | INGRAM_SHH_TARGETS_UP | 24 | -1.9658134 | 0.02981854 |
| 17 | DITTMER_PTHLH_TARGETS_DN | 16 | -1.932971 | 0.03479701 |
| 18 | MARTINEZ_RB1_TARGETS_UP | 125 | -1.9311029 | 0.03503092 |
| 19 | SANSOM_APC_TARGETS_REQUIRE_MYC | 35 | -1.9229536 | 0.03631372 |
| 20 | MARTINEZ_RB1_AND_TP53_TARGETS_UP | 105 | -1.9157431 | 0.03772556 |
| 21 | SHEN_SMARCA2_TARGETS_UP | 64 | -1.8830324 | 0.0409475 |
| 22 | MARTORIATI_MDM4_TARGETS_FETAL_LIVER_UP | 23 | -1.8816452 | 0.0411273 |
| 23 | GRABARCZYK_BCL11B_TARGETS_UP | 17 | -1.8777169 | 0.04166126 |
| 24 | GOZGIT_ESR1_TARGETS_UP | 35 | -1.8768103 | 0.04142939 |
| 25 | BEGUM_TARGETS_OF_PAX3_FOXO1_FUSION_UP | 15 | -1.8691893 | 0.0428513 |
| 26 | LIU_CMYB_TARGETS_UP | 32 | -1.8599355 | 0.04410038 |
| 27 | LIU_SOX4_TARGETS_UP | 27 | -1.8516853 | 0.04590253 |
| 28 | MARTINEZ_TP53_TARGETS_UP | 103 | -1.842061 | 0.04834863 |
| 29 | MARTORIATI_MDM4_TARGETS_NEUROEPITHELIUM_UP | 34 | -1.8389077 | 0.04845129 |
| 30 | PURBEY_TARGETS_OF_CTBP1_NOT_SATB1_UP | 46 | -1.8330786 | 0.04922288 |
| 31 | YANG_BCL3_TARGETS_UP | 58 | 2.107844 | 0.04891814 |

**Supplementary Table 7** GBM1 H3K27me3 GSEA gene sets containing target genes

| | NAME | SIZE | NES | QVAL |
|---|---|---|---|---|
| 40 | ACOSTA_PROLIFERATION_INDEPENDENT_MYC_TARGETS_UP | 27 | -2.24911 | 0.00570056 |
| 79 | BAE_BRCA1_TARGETS_UP | 28 | -1.9162892 | 0.0328236 |
| 48 | BASAKI_YBX1_TARGETS_DN | 140 | -2.1601255 | 0.00953125 |
| 77 | BASAKI_YBX1_TARGETS_UP | 90 | -1.92177 | 0.032053 |
| 34 | BENPORATH_MYC_MAX_TARGETS | 155 | -2.3689542 | 0.00263608 |
| 84 | BENPORATH_MYC_TARGETS_WITH_EBOX | 59 | -1.8642898 | 0.04217548 |
| 29 | BENPORATH_NANOG_TARGETS | 292 | -2.4994984 | 0.00093452 |
| 17 | BENPORATH_SOX2_TARGETS | 211 | -2.6754265 | 0.00022024 |
| 64 | BONCI_TARGETS_OF_MIR15A_AND_MIR16_1 | 37 | -2.0240557 | 0.01931548 |
| 15 | CHEN_HOXA5_TARGETS_9HR_UP | 61 | -2.698807 | 0.00015527 |
| 26 | CUI_TCF21_TARGETS_2_DN | 364 | -2.5718772 | 0.00054619 |
| 9 | DOUGLAS_BMI1_TARGETS_DN | 101 | -2.9422581 | 1.41E-05 |
| 56 | DOUGLAS_BMI1_TARGETS_UP | 200 | -2.0826068 | 0.0146461 |
| 52 | FEVR_CTNNB1_TARGETS_DN | 177 | -2.112302 | 0.01263329 |
| 38 | FISCHER_DIRECT_P53_TARGETS_META_ANALYSIS | 104 | -2.2723205 | 0.00497944 |
| 2 | FISCHER_DREAM_TARGETS | 211 | -3.457577 | 0 |
| 60 | FORTSCHEGGER_PHF8_TARGETS_DN | 259 | -2.0576117 | 0.01654953 |
| 27 | FUJII_YBX1_TARGETS_DN | 52 | -2.5349739 | 0.00069425 |
| 16 | GABRIELY_MIR21_TARGETS | 110 | -2.6804252 | 0.00021124 |
| 67 | GARCIA_TARGETS_OF_FLI1_AND_DAX1_DN | 45 | -1.9665704 | 0.02608781 |
| 5 | GARY_CD5_TARGETS_DN | 114 | -3.1042466 | 0 |
| 57 | GARY_CD5_TARGETS_UP | 131 | -2.0790005 | 0.01488459 |
| 24 | GAUSSMANN_MLL_AF4_FUSION_TARGETS_A_UP | 71 | -2.6079042 | 0.00039724 |
| 73 | GAUSSMANN_MLL_AF4_FUSION_TARGETS_E_UP | 46 | -1.9306456 | 0.03073017 |
| 80 | GEORGES_CELL_CYCLE_MIR192_TARGETS | 15 | -1.8986295 | 0.03589996 |
| 25 | GEORGES_TARGETS_OF_MIR192_AND_MIR215 | 259 | -2.580959 | 0.00048891 |
| 33 | GLASS_IGF2BP1_CLIP_TARGETS_KNOCKDOWN_DN | 36 | -2.380038 | 0.00246929 |
| 37 | GOBP_PROTEIN_TARGETING | 102 | -2.2959447 | 0.00422668 |
| 65 | GOBP_PROTEIN_TARGETING_TO_LYSOSOME | 16 | -2.0086527 | 0.02076599 |
| 46 | GOBP_PROTEIN_TARGETING_TO_VACUOLE | 23 | -2.1700191 | 0.00903752 |
| 81 | GOZGIT_ESR1_TARGETS_DN | 332 | -1.8830141 | 0.03869561 |
| 11 | HALLMARK_E2F_TARGETS | 53 | -2.9088047 | 1.21E-05 |
| 39 | HALLMARK_MYC_TARGETS_V1 | 36 | -2.2601438 | 0.00537506 |
| 63 | HAN_SATB1_TARGETS_DN | 141 | -2.0423954 | 0.01766531 |
| 32 | HELLER_HDAC_TARGETS_UP | 90 | -2.435981 | 0.00153838 |
| 30 | HINATA_NFKB_TARGETS_FIBROBLAST_UP | 41 | -2.4988525 | 0.00093904 |
| 86 | HINATA_NFKB_TARGETS_KERATINOCYTE_UP | 41 | -1.8534136 | 0.04440295 |
| 36 | HOOI_ST7_TARGETS_DN | 51 | -2.3408532 | 0.00317312 |
| 14 | HORIUCHI_WTAP_TARGETS_DN | 95 | -2.7460866 | 7.83E-05 |
| 90 | HORIUCHI_WTAP_TARGETS_UP | 120 | -1.8323499 | 0.04832166 |
| 76 | INGRAM_SHH_TARGETS_DN | 29 | -1.9223626 | 0.03195543 |

| | | | | |
|---|---|---|---|---|
| 62 | IVANOVSKA_MIR106B_TARGETS | 25 | -2.0476108 | 0.01731584 |
| 10 | JACKSON_DNMT1_TARGETS_UP | 35 | -2.9179108 | 1.29E-05 |
| 28 | JOHNSTONE_PARVB_TARGETS_2_DN | 110 | -2.5268078 | 0.00074562 |
| 3 | JOHNSTONE_PARVB_TARGETS_3_DN | 218 | -3.4222603 | 0 |
| 23 | JOHNSTONE_PARVB_TARGETS_3_UP | 156 | -2.6086023 | 0.00039898 |
| 47 | KIM_WT1_TARGETS_DN | 123 | -2.1656947 | 0.00931014 |
| 68 | KINSEY_TARGETS_OF_EWSR1_FLII_FUSION_DN | 118 | -1.9593173 | 0.02684986 |
| 8 | KINSEY_TARGETS_OF_EWSR1_FLII_FUSION_UP | 399 | -2.9666076 | 1.57E-05 |
| 21 | KOINUMA_TARGETS_OF_SMAD2_OR_SMAD3 | 300 | -2.611937 | 0.00037677 |
| 69 | KRIEG_KDM3A_TARGETS_NOT_HYPOXIA | 75 | -1.9437785 | 0.02889289 |
| 12 | LEE_BMP2_TARGETS_DN | 266 | -2.8425908 | 2.01E-05 |
| 70 | LIN_NPAS4_TARGETS_UP | 60 | -1.9365249 | 0.0299572 |
| 18 | LINSLEY_MIR16_TARGETS | 64 | -2.6447592 | 0.00026672 |
| 22 | LIU_SOX4_TARGETS_DN | 74 | -2.6094892 | 0.00039282 |
| 6 | LOPEZ_MBD_TARGETS | 283 | -3.0958116 | 0 |
| 20 | LU_EZH2_TARGETS_DN | 149 | -2.6179912 | 0.00035373 |
| 51 | MARTINEZ_RB1_AND_TP53_TARGETS_UP | 223 | -2.1128986 | 0.01258908 |
| 75 | MARTINEZ_RB1_TARGETS_DN | 195 | -1.9249184 | 0.03147958 |
| 85 | MARTINEZ_RB1_TARGETS_UP | 256 | -1.8573843 | 0.04355692 |
| 58 | MARTINEZ_TP53_TARGETS_UP | 225 | -2.0719457 | 0.0153684 |
| 19 | MARTORIATI_MDM4_TARGETS_FETAL_LIVER_DN | 185 | -2.6281588 | 0.00033726 |
| 53 | MOHANKUMAR_HOXA1_TARGETS_UP | 127 | -2.0907423 | 0.01410227 |
| 87 | MOLENAAR_TARGETS_OF_CCND1_AND_CDK4_UP | 24 | -1.852264 | 0.04460762 |
| 1 | NUYTTEN_EZH2_TARGETS_DN | 277 | -3.4639895 | 0 |
| 83 | NUYTTEN_EZH2_TARGETS_UP | 401 | -1.8744963 | 0.04021444 |
| 59 | NUYTTEN_NIPP1_TARGETS_DN | 261 | -2.0631604 | 0.01608984 |
| 54 | NUYTTEN_NIPP1_TARGETS_UP | 272 | -2.083825 | 0.01459737 |
| 61 | ODONNELL_TFRC_TARGETS_DN | 41 | -2.0552852 | 0.01679982 |
| 41 | RODRIGUES_DCC_TARGETS_DN | 43 | -2.2292986 | 0.00647713 |
| 42 | SANSOM_APC_MYC_TARGETS | 72 | -2.225965 | 0.00658031 |
| 45 | SCHLOSSER_MYC_TARGETS_REPRESSED_BY_SERUM | 48 | -2.180616 | 0.00849909 |
| 72 | SCIBETTA_KDM5B_TARGETS_DN | 20 | -1.9311659 | 0.03065723 |
| 31 | SENESE_HDAC1_TARGETS_UP | 182 | -2.4443629 | 0.00145532 |
| 74 | SENESE_HDAC3_TARGETS_UP | 198 | -1.927394 | 0.03120848 |
| 7 | SHEN_SMARCA2_TARGETS_UP | 121 | -3.0267565 | 0 |
| 44 | STEIN_ESRRA_TARGETS | 154 | -2.1809378 | 0.00849654 |
| 50 | STEIN_ESRRA_TARGETS_DN | 26 | -2.1354134 | 0.01116521 |
| 82 | STEIN_ESRRA_TARGETS_UP | 118 | -1.8774377 | 0.03963048 |
| 4 | TOYOTA_TARGETS_OF_MIR34B_AND_MIR34C | 126 | -3.3378465 | 0 |
| 66 | UDAYAKUMAR_MED1_TARGETS_UP | 37 | -1.9743384 | 0.02508564 |
| 88 | VANDESLUIS_COMMD1_TARGETS_GROUP_3_DN | 17 | -1.8517543 | 0.04462018 |
| 71 | WANG_LMO4_TARGETS_DN | 105 | -1.9344578 | 0.03026319 |
| 35 | WANG_LMO4_TARGETS_UP | 119 | -2.3613951 | 0.00276263 |

| | | | | |
|---|---|---|---|---|
| 49 | WANG_SMARCE1_TARGETS_DN | 131 | -2.1444209 | 0.01055117 |
| 13 | WEI_MYCN_TARGETS_WITH_E_BOX | 227 | -2.7623253 | 4.91E-05 |
| 55 | WELCSH_BRCA1_TARGETS_DN | 35 | -2.0837376 | 0.01458793 |
| 89 | YAMAZAKI_TCEB3_TARGETS_UP | 65 | -1.8323545 | 0.04835339 |
| 78 | YANG_BCL3_TARGETS_UP | 136 | -1.9168502 | 0.032766 |
| 43 | YOSHIMURA_MAPK8_TARGETS_DN | 132 | -2.1840134 | 0.00832685 |
